## Supplementary Text for "Resolution of the curse of dimensionality in single-cell RNA sequencing data analysis"

### Mathematical background
### for
### resolution of the curse of dimensionality in single-cell RNA sequencing data analysis

Yusuke Imoto[1], Tomonori Nakamura[1,2,3], Emerson G. Escolar[4,5], Michio Yoshiwaki[5], Yoji Kojima[1,2,6], Yukihiro Yabuta[1,2], Yoshitaka Katou[2], Takuya Yamamoto[1,5,6], Yasuaki Hiraoka[1,5,7], and Mitinori Saitou[1,2,6].

Contents


1. Curse of dimensionality 2
1.1. Curse of dimensionality 1: Loss of closeness 3
1.2. Curse of dimensionality 2: Inconsistency of statistics 4
2. Existing statistical approaches for the curse of dimensionality 6
2.1. Noise reduction methodology 6
2.2. Cross-data matrix methodology 7
2.3. Comparison of NRM and CDM 7
3. RECODE 8
3.1. Method 8
3.2. Theory and parameter estimation 11
3.3. Computation 14
3.4. Verification 15
4. RECODE for single-cell RNA-sequencing data 18
4.1. Statistical theory and method 18
4.2. Computation 22
4.3. Verification by scRNA-seq data 23
5. Extensions of RECODE 30
5.1. Fast algorithm 30
5.2. Applicability 31
5.3. Extension to single-cell epigenome data 32
5.4. Discussion of extension to future data 34
Appendix A. Mathematical notation 35
Appendix B. Proofs 35
References 39


This supplementary text was mainly written by Y. Imoto, E. G. Escolar, M. Yoshiwaki, and Y. Hiraoka.

### 1. Curse of dimensionality

We consider high-dimensional observed data containing experimental (technical) noise. Assume that we have $n$ samples (observations) with $d$ features, with the $j$th observation being represented by the $d$-dimensional real-valued column vector $x_j = (x_{1j}, \ldots, x_{dj})^{\mathrm{T}} \in \mathbb{R}^d$ consisting of measured values for each of those features. We denote the observed data matrix as $X = (x_1, \ldots, x_n) \in \mathbb{R}^{d \times n}$.

In general, we cannot determine the true values because of experimental errors in data sampling. We model the presence of noise by variables $e_{ij}$ for $i = 1, \ldots, d$ and $j = 1, \ldots, n$. Their values are defined as the differences between the observed and true values.

$$e_{ij} := x_{ij} - x_{ij}^{\mathtt{true}}.$$

Here, $x_{ij}^{\mathtt{true}}$ represents the true value corresponding to $x_{ij}$. Now, we assume that for each index of feature $i = 1, \ldots, d$, the random variables $x_{i1}, \ldots, x_{in}$ ($e_{i1}, \ldots, e_{in}$, respectively) are independent and identically distributed. We denote the $j$th true data vector and true data matrix as $x_j^{\mathtt{true}} = (x_{1j}^{\mathtt{true}}, \ldots, x_{dj}^{\mathtt{true}})^{\mathrm{T}} \in \mathbb{R}^d$ and $X^{\mathtt{true}} = (x_1^{\mathtt{true}}, \ldots, x_n^{\mathtt{true}}) \in \mathbb{R}^{d \times n}$, respectively. Thus, we similarly have noise vectors

$$e_j := x_j - x_j^{\mathtt{true}}$$

for $j = 1, \ldots, n$.

For example, when the observed data are the gene expression levels detected by single-cell RNA sequencing (scRNA-seq), the true data are the actual amounts of RNA in each single cell, and the sample size $n$ and dimension $d$ represent the number of single cells and genes, respectively (Fig ST1). In general, gene expression data can be considerably high-dimensional. For example, mammalian cells contain approximately 20,000 or more genes.

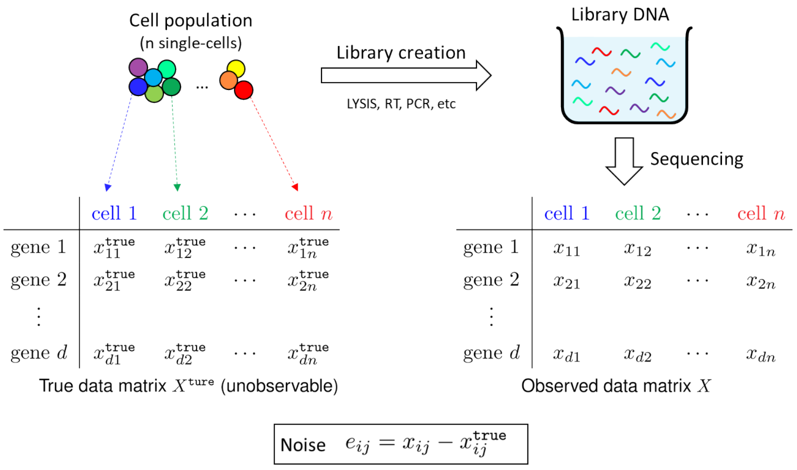


Figure ST1. Diagram of true data, observed data, and noise in scRNA-seq data.

When analyzing high-dimensional observed data, the "curse of dimensionality" (COD), encompassing many issues in this setting, must be addressed. In this study, we consider the problem of noise significantly affecting the computation of values, such as distances, if the dimension $d$ is extremely high. This occurs even if the noise $e$ for each coordinate is minimal. In particular, we identify two main problems: the *loss of closeness* (COD1) and *inconsistency of statistics* (COD2).

1.1. **Curse of dimensionality 1: Loss of closeness.** We present several mathematical results that characterize the COD by affecting the computation of distances between observations. First, we obtain the following proposition.

**Proposition 1.1** (**Curse of dimensionality 1-i**). *Assume that the noise vectors $e_1, e_2, \ldots, e_n$ are independent and identically distributed (i.i.d.) $d$-dimensional random variables, each with mean vector $0$ and covariance matrix $\sigma^2 I_d$, where $\sigma^2$ is a positive real number and $I_d$ is the $d \times d$ identity matrix. Moreover, let $x_j^{\mathtt{true}}$ and $e_{j'}$ be independent of each $j, j' = 1, \ldots, n$. Then, the expected value of the squared Euclidean distance between $x_j$ and $x_{j'}$ for distinct $j, j' \in \{1, \ldots, n\}$ satisfies*

$$\mathrm{E}(\|x_j - x_{j'}\|^2) = \mathrm{E}(\|x_j^{\mathtt{true}} - x_{j'}^{\mathtt{true}}\|^2) + 2\sigma^2 d,$$

*where $\|\cdot\|$ is the Euclidean norm.*

We provide a proof in Appendix B. In the following, we present an interpretation of the result. If $\|x_j^{\mathtt{true}} - x_{j'}^{\mathtt{true}}\|$ is sufficiently large ($\|x_j^{\mathtt{true}} - x_{j'}^{\mathtt{true}}\| \gg \sigma\sqrt{2d}$), then we can regard the distance between the observed data $x_j$ and $x_{j'}$ as a reasonable approximation of the true distance ($\|x_j - x_{j'}\| \approx \|x_j^{\mathtt{true}} - x_{j'}^{\mathtt{true}}\|$). In contrast, if $x_j^{\mathtt{true}}$ and $x_{j'}^{\mathtt{true}}$ are sufficiently close ($\|x_j^{\mathtt{true}} - x_{j'}^{\mathtt{true}}\| \ll \sigma\sqrt{2d}$), then the distance information is drowned out by the noise term $\sigma\sqrt{2d}$.

Proposition 1.1 assumes independence between $x_j^{\mathtt{true}}$ and $e_{j'}$ for the sake of simplicity. In the following, we do not assume that condition. We present another result related to the COD. We denote the $d$-dimensional multivariate Gaussian distribution with mean $\mu$ and the covariance matrix $\Sigma$ by $N(\mu, \Sigma)$.

**Proposition 1.2** (**Curse of dimensionality 1-ii** [8, Section 2]). *Assume that $e_j \sim N(0, I_d)$ are i.i.d. Then, for a distinct $j, j' \in \{1, \ldots, n\}$ such that $x_j^{\mathtt{true}} = x_{j'}^{\mathtt{true}}$,*

$$\|x_j - x_{j'}\| = \sqrt{2d} + O_{\mathtt{p}}(1) \qquad \text{as } d \to \infty.$$

Here, the notation $Y_d = O_{\mathtt{p}}(a_d)$ denotes the stochastic boundedness of $Y_d/a_d$ as $d \to \infty$ (see Appendix A). We consider equal true data $x_j^{\mathtt{true}} = x_{j'}^{\mathtt{true}}$; therefore, the true distance is 0. However, the observed distance scales as $\sqrt{2d}$; thus, it can be far from 0 for high dimensions.

In the case of noise distributions that are not Gaussian distributions, Yata and Aoshima [4] reported the following result, which is an extension of Proposition 1.2.

**Proposition 1.3** (**Curse of dimensionality 1-iii** [4]). *Let $x_1, \ldots, x_n$ be i.i.d. $d$-dimensional random variables with mean $\mu$ and covariance matrix $S$. If the conditions*

$$\frac{\mathrm{tr}(S^2)}{\mathrm{tr}(S)^2} \to 0 \qquad \text{as } d \to \infty \tag{1}$$

*and*

$$\frac{\mathrm{Var}(\|x_j - \mu\|^2)}{\mathrm{tr}(S)^2} \to 0 \qquad \text{as } d \to \infty \tag{2}$$

*are satisfied, then for $j, j' = 1, \ldots, n$ with $j \neq j'$, it holds that*

$$\|x_j - x_{j'}\| = \sqrt{2\mathrm{tr}(S)} + o_{\mathrm{p}}(\sqrt{\mathrm{tr}(S)}) \qquad \textit{as } d \to \infty.$$

Here, the notation $Y_d = o_{\mathrm{p}}(a_d)$ denotes the convergence to zero in the probability of $Y_d/a_d$ as $d \to \infty$ (see Appendix A). We summarize its proof [4] in Appendix B. In this setting, the observed distance scales as $\sqrt{2\mathrm{tr}(S)}$. Note that $\mathrm{tr}(S)$ is the sum of the variances of the components in the $d$-dimensional distribution, and this increases with $d$ assuming each component has some constant variance (noise). The results show that the Euclidean distances between observations in a high-dimensional space can contain large noise terms despite the small size of the noise in each coordinate. Many data analysis methods rely on the computation of the distances between points. Hence, if this issue is not addressed when dealing with high-dimensional data, the accuracy of the results may be questionable. However, although principal component analysis (PCA) does not require any distance calculations, it is affected by another aspect of the COD, as discussed in the next section.

1.2. **Curse of dimensionality 2: Inconsistency of statistics.** To explain the inconsistency of statistics arising from the COD, we first provide a quick review of PCA, as the inconsistency is closely related to the eigenvalues in PCA.

PCA is a multivariate analysis method that identifies a new coordinate system corresponding to the directions of the maximal variances of the data. The data can then be expressed in terms of the new coordinate system.

The underlying theory of PCA can be explained as follows. First, recall that the sample covariance matrix of $X$ is given as

$$S_X := \frac{1}{n-1}(X - \overline{X})(X - \overline{X})^{\mathrm{T}},$$

where $\overline{X} := XP$ and $P$ is an $n \times n$ matrix such that all the components are $1/n$. That is, $\overline{X}$ consists of the means of the rows of $X$. Then, we solve the eigenvalue problem $S_X u = \lambda u$ by using the following solutions:

$$S_X u_{X,i} = \lambda_{X,i} u_{X,i} \qquad \text{for } i = 1, \ldots, d. \tag{3}$$

where $\lambda_{X,i}$ are the eigenvalues and $u_{X,i}$ are the eigenvectors that are chosen such that $\lambda_{X,1} \geq \cdots \geq \lambda_{X,d}$ and that $U_X = (u_{X,1}, \ldots, u_{X,d})$ is an orthogonal matrix. Note that $\lambda_{X,i} = 0$ for $i = d^{\mathtt{PCA}} + 1, \ldots, d$, where $d^{\mathtt{PCA}}$ is the maximum dimension of the PCA-transformed data, that is, $d^{\mathtt{PCA}} := \min\{n-1, d\}$. The existence of such a solution is based on the basic linear algebra of symmetric matrices.

The transformed data matrix $X^{\mathtt{PCA}}$ in the PCA coordinate system are given as

$$X^{\mathtt{PCA}} := U_X^{\mathrm{T}}(X - \overline{X}).$$

We call projection $U_X^{\mathrm{T}}(\,\cdot\, - \overline{X})$ *PCA projection* for data matrix $X$. Let us consider the sample covariance matrix $S_{X^{\mathtt{PCA}}}$. Noting that $\overline{X^{\mathtt{PCA}}} = X^{\mathtt{PCA}}P = U_X^{\mathrm{T}}(X - XP)P = 0$ because $P^2 = P$, we obtain

$$S_{X^{\mathtt{PCA}}} = \frac{1}{n-1} X^{\mathtt{PCA}} {X^{\mathtt{PCA}}}^{\mathrm{T}} = U_X^{\mathrm{T}} S_X U_X = \Lambda,$$

where $\Lambda$ is a diagonal matrix with diagonal entries $\lambda_{X,1}, \ldots, \lambda_{X,d}$. That is, the eigenvalue $\lambda_{X,i}$ corresponds to the variance of the data in the $i$th principal component.

However, in the case of high-dimensional and low-sample-size data, these eigenvalues (and thus, the corresponding variances) may not converge to the true eigenvalues (true variances). [22, Theorem 3.1 and Theorem 3.3] (see Proposition 1.4). In particular, the explained variances in PCA may be inaccurate. Another perspective on this problem comes from an application of Chebyshev's inequality. For a random variable $x$ with mean $\mu$ and variance $\sigma^2$, the values that $x$ is likely to take can be characterized by Chebyshev's inequality.

$$\Pr(|x-\mu| \geq \eta\sigma) \leq \frac{1}{\eta^2} \qquad \text{for all } \eta > 0,$$

where $\Pr(\cdot)$ represents the probability. Chebyshev's inequality can be interpreted as a confidence interval based only on the mean and variance. Thus, the confidence intervals obtained from Chebyshev's inequality[1] may not reflect the true data space.

Let us review the aforementioned nonconvergence results. Hereafter in this section, we consider that the data $x_j$ $(j = 1, \ldots, n)$ are i.i.d. $d$-dimensional random variables with mean $\mu$ and covariance matrix $S$. We also assume that the true eigenvalues $\lambda_i$ $(i = 1, \ldots, d)$ of the covariance matrix $S$ are represented by the generalized spike model

$$\lambda_i = \begin{cases} c_i d^{\alpha_i}, & \text{for } i = 1, \ldots, m, \\ c_i, & \text{for } i = m+1, \ldots, d \end{cases}$$

with constants $c_i \geq 0$ for $i = 1, \ldots, d$ and $\alpha_1 \geq \cdots \geq \alpha_m > 0$ as the parameters. Here, we note that for $d \leq n$, the eigenvalue $\lambda_{X,i}$ is known to converge in probability to the true eigenvalue $\lambda_i$ as the sample size increases to infinity $(n \to \infty)$ [1]. However, this is not always true for $d > n$, which is considered high-dimensional and low-sample-size data.

Let $\zeta_j := \Lambda^{-1/2} U_X^{\mathrm{T}}(x_j - \mu)$ for each $j = 1, 2, \ldots, n$, where $\Lambda^{-1/2}$ is the diagonal matrix with diagonal entries $\lambda_1^{-1/2}, \ldots, \lambda_d^{-1/2}$. We consider the following condition:

(c0) The components $\zeta_{1j}, \ldots, \zeta_{dj}$ of $\zeta_j$ are independent for each $j = 1, \ldots, n$.

For example, if the data matrix $X$ follow a Gaussian distribution, then condition (c0) is satisfied. In general, the following statement holds.

**Proposition 1.4** (**Curse of dimensionality 2-i** [19, Corollary 1], [22, Theorem 3.1 and Theorem 3.3])**.** *Assume that condition* (c0) *holds. For $i = 1, 2, \ldots, m$ such that $\alpha_i > 0$, under the condition*

(c1) $$d \to \infty \text{ and } n \to \infty \text{ for } i \text{ such that } \alpha_i > 1$$

*or the condition*

(c2) $$d \to \infty \text{ and } d^{1-\alpha_i}/n \to 0 \text{ for } i \text{ such that } \alpha_i \in (0, 1],$$

*the convergence in probability*

$$\lambda_{X,i} \xrightarrow{\mathrm{p}} \lambda_i$$

*holds. Moreover, if $d^{1-\alpha_i}/n \not\to 0$ in condition* (c2)*, then*

$$\lambda_{X,i} \overset{\mathrm{p}}{\not\to} \lambda_i.$$

[1] In practice, Chebyshev's equality should be modified to use the sample mean and sample variance [15]. However, this does not resolve the nonconvergence of the variance.

Although the proof appears in the aforementioned references, we also provide an outline of the proof in Appendix B. Next, we consider cases in which condition (c0) is not satisfied, such as when the data are taken from a $t$-distribution. Without assuming condition (c0), the following holds.

**Proposition 1.5** (**Curse of dimensionality 2-ii** [19, Theorem 1])**.** *For* $i = 1, 2, \ldots, m$ *such that* $\alpha_i > 0$*, under condition* (c1) *or the condition*

(c3) $$d \to \infty \text{ and } d^{2-2\alpha_i}/n \to 0 \text{ for } i \text{ such that } \alpha_i \in (0, 1],$$

*the convergence in probability*

$$\lambda_{X,i} \xrightarrow{\mathrm{P}} \lambda_i$$

*holds.*

The difference between Proposition 1.4 and Proposition 1.5 is the exponent under conditions (c2) and (c3). Condition (c2) is weaker than condition (c3). From the propositions, as $d$ tends to infinity, even increasing the sample size $n$ to infinity cannot guarantee that the eigenvalues will converge to the true eigenvalues owing to the effects of noise. We illustrate the regions of convergence given by Proposition 1.4 and Proposition 1.5 in Fig ST2 (a) and (b), respectively.

#### 2. Existing statistical approaches for the curse of dimensionality

In this section, we review the existing statistical approaches established by Aoshima, Yata, and others [2, 19, 20, 21] for COD2 (inconsistency of statistics). Although the authors reported mathematical results for eigenvectors and PC scores, we only focus on the eigenvalues of the covariance matrix of data in this section. In particular, we review the noise reduction methodology [21] in Section 2.1 and cross-data matrix methodology [20] in Section 2.2. These methodologies are necessary to develop RECODE in the following section.

##### 2.1. Noise reduction methodology.

Yata and Aoshima [21] developed the *noise reduction methodology* (NRM) that modifies the eigenvalues of the covariance matrix of data in order to solve COD2 under condition (c0). The eigenvalues after the modification by the NRM are given as follows:

**Noise reduction methodology** [21]: The NRM defines the modified eigenvalues $\widetilde{\lambda}^{\mathtt{NRM}}_{X,i}$ $(i = 1, 2, \ldots, d)$ as

(4) $$\widetilde{\lambda}^{\mathtt{NRM}}_{X,i} := \begin{cases} \lambda_{X,i} - \dfrac{1}{d^{\mathtt{PCA}} - i + 1} \displaystyle\sum_{j=i+1}^{d^{\mathtt{PCA}}} \lambda_{X,j}, & i = 1, 2, \ldots, d^{\mathtt{PCA}}, \\ 0, & \text{otherwise.} \end{cases}$$

The key point of modifying the eigenvalues is given by the following result [21], which corresponds to an improvement in Proposition 1.4 because the conditions for convergence are weakened.

**Proposition 2.1** ([21, Theorem 3])**.** *Assume condition* (c0)*. For* $i = 1, 2, \ldots, m$*, under the conditions*

(c1′) $$d \to \infty \text{ and } n \to \infty \text{ for } i \text{ such that } \alpha_i > 1/2.$$

*or*

(c2′) $\quad d \to \infty$ *and* $d^{1-2\alpha_i}/n \to 0$ *for* $i$ *such that* $\alpha_i \in (0, 1/2]$,

*the convergence in probability*

$$\widetilde{\lambda}^{\mathtt{NRM}}_{X,i} \xrightarrow{\mathtt{P}} \lambda_i$$

*holds.*

2.2. **Cross-data matrix methodology.** In cases in which the data do not satisfy condition (c0) (e.g., data generated from a multidimensional $t$-distribution), Proposition 2.1 cannot be used. Accordingly, the NRM does not ensure the resolution of COD2. In such a setting, Yata and Aoshima proposed another method called the *cross-data matrix methodology* (CDM) [20].

**Cross-data matrix method** [20]: We divide data matrix $X = (x_1, \dots, x_n)$ into two disjoint datasets $X^{(1)} = (x^{(1)}_1, \dots, x^{(1)}_{n_1})$ and $X^{(2)} = (x^{(2)}_1, \dots, x^{(2)}_{n_2})$ such that $n_1 \geq n_2$ and $|n_1 - n_2| \leq 1$. The cross-data matrix $S^{\mathtt{CDM}}$ is defined as

$$S^{\mathtt{CDM}} = \frac{(X^{(1)} - \overline{X}^{(1)})^{\mathrm{T}}(X^{(2)} - \overline{X}^{(2)})}{\sqrt{(n_1-1)(n_2-1)}},$$

where $\overline{X}^{(k)} = X^{(k)}P$ for $k = 1, 2$. Then, the CDM defines the modified eigenvalues $\widetilde{\lambda}^{\mathtt{CDM}}_{X,i}$ $(i = 1, 2, \dots, n_2 - 1)$ by the singular value decomposition of the cross-data matrix:

$$S^{\mathtt{CDM}} = \sum_{i=1}^{n_2-1} \widetilde{\lambda}^{\mathtt{CDM}}_{X,i} u^{(1)}_i (u^{(2)}_i)^{\mathrm{T}} \tag{5}$$

with $\widetilde{\lambda}^{\mathtt{CDM}}_{X,1} \geq \cdots \geq \widetilde{\lambda}^{\mathtt{CDM}}_{X,n_2-1}$.

Then, the following holds for the CDM-modified eigenvalues [20, Theorem 1]:

**Proposition 2.2.** *For* $j = 1, 2, \dots, m$, *under condition* (c1′), *or the condition*

(c3′) $\quad d \to \infty$ *and* $d^{2-2\alpha_i}/n \to 0$ *for* $i$ *such that* $\alpha_i \in (0, 1/2]$,

*the convergence in probability*

$$\widetilde{\lambda}^{\mathtt{CDM}}_{X,i} \xrightarrow{\mathtt{P}} \lambda_i$$

*holds.*

Similar to Proposition 2.1, Proposition 2.2 improves Proposition 1.5 because it requires a weaker condition for convergence.

2.3. **Comparison of NRM and CDM.** Finally, in Fig ST2, we illustrate the differences between the conditions required for convergence in Proposition 1.4, Proposition 1.5, Proposition 2.1, and Proposition 2.2 when the sample size $n$ and dimension $d$ are related as $n = cd^{\gamma}$.

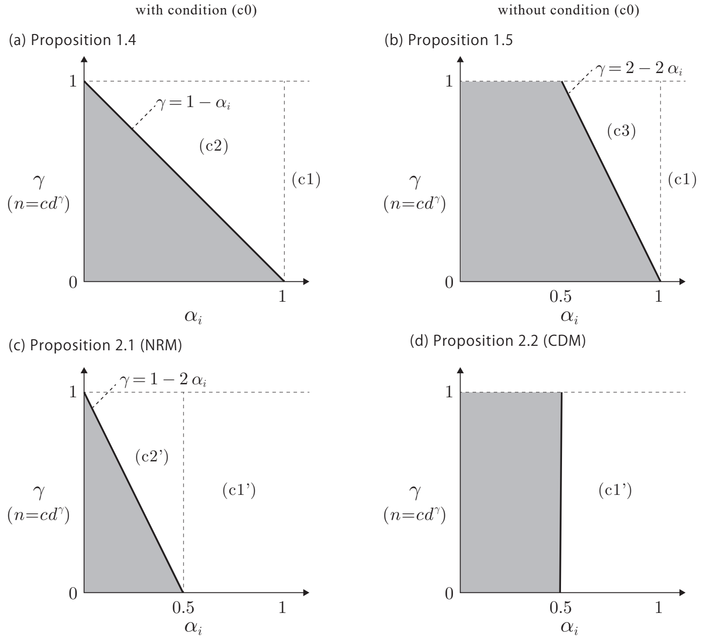


Figure ST2. Regions corresponding to conditions (c1), (c2), (c3), (c1′), and (c2′) that appear in Proposition 1.4, Proposition 1.5, Proposition 2.1, and Proposition 2.2 in the case of $n = cd^{\gamma}$, where $c$ is a positive constant independent of $n$ and $d$. The gray areas indicate the regions of non- or unknown convergence in probability. The schematic views of (a) and (c) were originally presented in [3] (see also [2]).

#### 3. RECODE

This section discusses our approach to resolving COD1 (loss of closeness) and COD2 (inconsistency of statistics).

**3.1. Method.** First, we develop a method that modifies the observed data to resolve COD2. In other words, the eigenvalues of the modified observed data can be obtained to match the modified eigenvalues in Section 2.

For a high-dimensional observed data matrix $X \in \mathbb{R}^{d \times n}$ and given any nonnegative real numbers $\widetilde{\lambda}_{X,i}$ , we construct a modified data matrix $\widetilde{X} \in \mathbb{R}^{d \times n}$ satisfying the following requirements:

$$\mu_{\widetilde{X}} = \mu_X, \tag{6}$$

$$S_{\widetilde{X}} u_{X,i} = \lambda_{\widetilde{X},i} u_{X,i} \quad \text{for } i = 1, 2, \ldots, d, \tag{7}$$

$$\lambda_{\widetilde{X},i} = \widetilde{\lambda}_{X,i} \quad \text{for } i = 1, 2, \ldots, d. \tag{8}$$

Here, $\mu_X$ and $\mu_{\widetilde{X}}$ are the mean vectors of $X$ and $\widetilde{X}$, respectively. Moreover, $S_{\widetilde{X}}$ is the sample covariance matrix of $\widetilde{X}$.

The first requirement (6) simply requires the same central position for $X$ and $\tilde{X}$. The second requirement (7) means that the original eigenvectors also serve as eigenvectors for the modified data. That is, the structure of the data extracted by PCA is preserved. The third requirement (8) means that the eigenvalues of the sample covariance matrix of $\widetilde{X}$ coincide with the chosen eigenvalues $\widetilde{\lambda}_{X,i}$. We set the eigenvalues $\widetilde{\lambda}_{X,i}$ $(i = 1, 2, \ldots, d)$ as ones $\widetilde{\lambda}^{\texttt{NRM}}_{X,i}$ defined by the NRM in Eq. (4). Alternatively, if the data are not assumed to satisfy condition (c0), then we can use the eigenvalues $\widetilde{\lambda}^{\texttt{CDM}}_{X,i}$ defined by the CDM in Eq. (5).

We need the following lemma for the construction (see Appendix B for the proof).

**Lemma 3.1.** *For a diagonal matrix $L \in \mathbb{R}^{d\times d}$, let*

$$Y \coloneqq U_X L U_X^{\mathrm{T}}(X - \overline{X}) + \overline{X}. \tag{9}$$

*Then, the mean $\mu_Y$ and covariance matrix $S_Y$ of $Y$ satisfy*

$$\mu_Y = \mu_X$$

*and*

$$S_Y u_{X,i} = \lambda_{X,i} L_{ii}^2 u_{X,i} \quad \text{for } i = 1, 2, \ldots, d,$$

*respectively.*

Lemma 3.1 shows that the transformation in Eq. (9) can arbitrarily modify the eigenvalues of the covariance matrix of the data while satisfying the requirements (6) and (7). Let $\widetilde{\Lambda}_X^{1/2}$ and $\Lambda_X^{-1/2}$ be $d \times d$ diagonal matrices with diagonal entries $\widetilde{\lambda}_{X,1}^{1/2}, \ldots, \widetilde{\lambda}_{X,d}^{1/2}$ and $\lambda_{X,1}^{-1/2}, \ldots, \lambda_{X,d}^{-1/2}$, respectively, where we set $(\Lambda_X^{-1/2})_{ii} = 0$ when $\lambda_{X,i} = 0$. Then, replacing $L$ in Lemma 3.1 with $\widetilde{\Lambda}_X^{1/2}\Lambda_X^{-1/2}$ ($L_{ii}^2 = \widetilde{\lambda}_{X,i}/\lambda_{X,i}$ for $i$ such that $\lambda_{X,i} \neq 0$), we define a data matrix $\widetilde{X}$ as

$$\widetilde{X} \coloneqq U_X \widetilde{\Lambda}_X^{1/2} \Lambda_X^{-1/2} U_X^{\mathrm{T}}(X - \overline{X}) + \overline{X}. \tag{10}$$

Then, from Lemma 3.1, the following theorem immediately holds.

**Theorem 3.2.** *For a data matrix $X \in \mathbb{R}^{d\times n}$ and given any nonnegative real number $\widetilde{\lambda}_{X,i}$, the data matrix $\widetilde{X}$, as defined in Eq. (10), satisfies the requirements (6)–(8).*

Suppose we compute the modified data matrix $\widetilde{X}$ using the eigenvalues defined by the NRM in Eq. (4) or the CDM in Eq. (5). Then, under the conditions of Proposition 2.1 or Proposition 2.2, the modified eigenvalues converge to the true eigenvalues $\lambda_i$. From Theorem 3.2, the eigenvalues of the modified data matrix $\widetilde{X}$ are exactly equal to the modified eigenvalues. That is, COD2 is resolved.

However, the modified data could still be affected by COD1. To comprehensively resolve COD1 and COD2, we propose the following noise reduction method called *RECODE* (resolution of the curse of dimensionality).

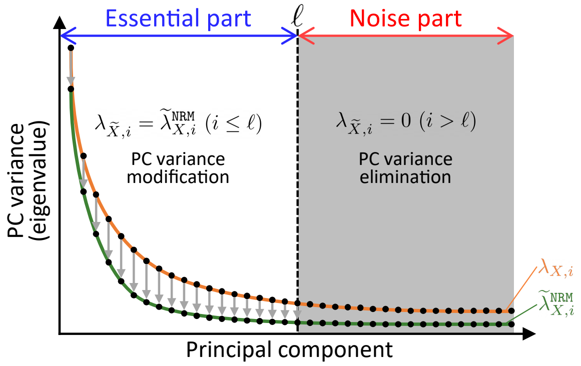


Figure ST3. Illustration of the eigenvalue transformation by RECODE.

**RECODE**

For a data matrix $X$ and parameter $\ell \in \{1, 2, \ldots, d\}$, RECODE defines the modified data matrix

$$\widetilde{X}_\ell := U_X \widetilde{\Lambda}^{1/2}_{X,\ell} \Lambda_X^{-1/2} U_X^{\mathrm{T}} (X - \overline{X}) + \overline{X}, \tag{11}$$

where $\widetilde{\Lambda}^{1/2}_{X,\ell} = \mathrm{diag}(\widetilde{\lambda}^{1/2}_{X,1}, \ldots, \widetilde{\lambda}^{1/2}_{X,d})$ and

$$\widetilde{\lambda}_{X,i} := \begin{cases} \widetilde{\lambda}^{\mathtt{NRM}}_{X,i}, & i \leq \ell \text{ (PC variance modification)}, \\ 0, & i > \ell \text{ (PC variance elimination)}. \end{cases}$$

Using a proof similar to that for Theorem 3.2, we present the following theorem.

**Theorem 3.3.** *The RECODE-modified data matrix $\widetilde{X}_\ell$ satisfies the requirement (6)–(8). In particular, $\lambda_{\widetilde{X}_\ell, i} = 0$ for $i = \ell + 1, \ldots, d$.*

RECODE regards the first $\ell$ principal components (PCs) as the essential part and the other PCs as the noise part. Thus, RECODE transforms the PC variances (eigenvalues) by the NRM (4) for the essential part (PC variance modification) and them to zero to eliminate the noise effect for the noise part (PC variance elimination) (Fig ST3).

We provide the following result, which explains the modification made by RECODE from another perspective.

**Proposition 3.4.** *Let $X - \overline{X} = U_X \Sigma V_X^{\mathrm{T}}$ be the singular value decomposition of $X - \overline{X}$. Then, the RECODE-modified data matrix $\widetilde{X}_\ell$ can be represented as*

$$\widetilde{X}_\ell - \overline{X} = U_X \widetilde{\Sigma} V_X^{\mathrm{T}}.$$

*Here, $\widetilde{\Sigma}$ is the $d \times n$ rectangular diagonal matrix with diagonal entries given by*

$$\widetilde{\Sigma}_{ii} = \begin{cases} \sqrt{(n-1)\widetilde{\lambda}_{X,i}}, & i = 1, \ldots, \ell, \\ 0, & i = \ell + 1, \ldots, \min\{n - 1, d\}. \end{cases}$$

*In particular, when $\widetilde{\lambda}_{X,i}$ is given by the NRM (4),*

$$\widetilde{\Sigma}_{ii} = \begin{cases} \sqrt{\Sigma_{ii}^2 - \dfrac{1}{d^{\mathtt{PCA}} - i + 1} \displaystyle\sum_{j=i+1}^{d^{\mathtt{PCA}}} \Sigma_{jj}^2}, & i = 1, \ldots, \ell, \\ 0, & \text{otherwise}. \end{cases}$$

Proposition 3.4 can be proved using the fact that $\Sigma$ is the diagonal matrix $\mathrm{diag}(\sqrt{(n-1)\lambda_{X,1}}, \ldots, \sqrt{(n-1)\lambda_{X,d}})$. From Proposition 3.4, RECODE can be interpreted as a modification of the singular values of the centered data matrix $X - \overline{X}$ (Fig ST4).

In the next section, we present a procedure for estimating the parameter $\ell$ for the data and show that RECODE resolves COD1.

3.2. **Theory and parameter estimation.** In this section, we introduce theories of RECODE and a method for inferring parameter $\ell$. To this end, we assume the following condition:

(C1) There exists $m < d^{\mathtt{PCA}}$ such that the information of the true data matrix $X^{\mathtt{true}}$ is fully explained by the first $m$ PCs of the observed data, that is,

$$u_{X,i}^{\mathrm{T}}(X^{\mathtt{true}} - \overline{X^{\mathtt{true}}}) = 0 \text{ for } i = m+1, \ldots, d.$$

Here, we note that when $X^{\mathtt{true}}$ is sampled from a finite-dimensional manifold, condition (C1) is satisfied by increasing the sample size $n$. Under condition (C1), the information of the true data can be fully captured using at most the first $m$ PCs of the observed data, that is, a PCA space with a dimension of at most $m$. Therefore, the optimal value for the parameter $\ell$ of RECODE is the minimum $m$ for condition (C1). Moreover, we obtain the following theorem, which can be seen as a resolution, via RECODE, of the COD, as shown in Proposition 1.1.

**Theorem 3.5.** *For a fixed data matrix $X \in \mathbb{R}^{d\times n}$, let $U_X$ and $\Lambda_X$ be defined from the eigenvalue decomposition $S_X = U_X \Lambda_X U_X^{\mathrm{T}}$ of its covariance matrix $S_X$. Let $\widetilde{\lambda}_{X,i}$ $(i = 1, 2, \ldots, d)$ be modified eigenvalues such that $\widetilde{\lambda}_{X,i} \leq \lambda_{X,i}$. Assume that the noise $e_j = (e_{1j}, \ldots, e_{dj})^{\mathrm{T}}$ $(j = 1, \ldots, n)$ are i.i.d. random vectors with mean $0$ and covariance matrix $\sigma^2 I_d$ and that $x_j^{\mathtt{true}}$ and $e_j$ are independent for $j = 1, \ldots, n$. Then, for distinct $j, j' \in \{1, \ldots, n\}$ with the random variables $x_j = x_j^{\mathtt{true}} + e_j$ and $x_{j'} = x_{j'}^{\mathtt{true}} + e_{j'}$ satisfying the condition that for some $m < \min\{n-1, d\}$,*

$$u_{X,i}^{\mathrm{T}}(x_k^{\mathtt{true}} - \mu_{X^{\mathtt{true}}}) = 0 \quad \textit{for } i = m+1, \ldots, \min\{n-1, d\},\ k \in \{j, j'\}, \tag{12}$$

*the inequality*

$$\mathrm{E}(\|\widetilde{x}_j - \widetilde{x}_{j'}\|^2) \leq \mathrm{E}(\|x_j^{\mathtt{true}} - x_{j'}^{\mathtt{true}}\|^2) + 2\sigma^2 m \tag{13}$$

*holds, where $\widetilde{x}_j$ and $\widetilde{x}_{j'}$ are defined by*

$$\widetilde{x}_k := U_X \widetilde{\Lambda}_{X,m}^{1/2} \Lambda_X^{-1/2} U_X^{\mathrm{T}}(x_k - \mu_X) + \mu_X, \quad k \in \{j, j'\}. \tag{14}$$

The proof is provided in Appendix B. Note that for this theorem, the random variables $x_j$ and $x_{j'}$ are not columns from the fixed data matrix $X$, which is constant. Condition (12) corresponds to condition (C1), and the transformation (14) corresponds to transforming $x_k$ using RECODE (11) based on the fixed data matrix $X$. Moreover, the condition $\widetilde{\lambda}_{X,i} \leq \lambda_{X,i}$ is satisfied if we use the modified eigenvalues provided by the NRM in Eq. (4).

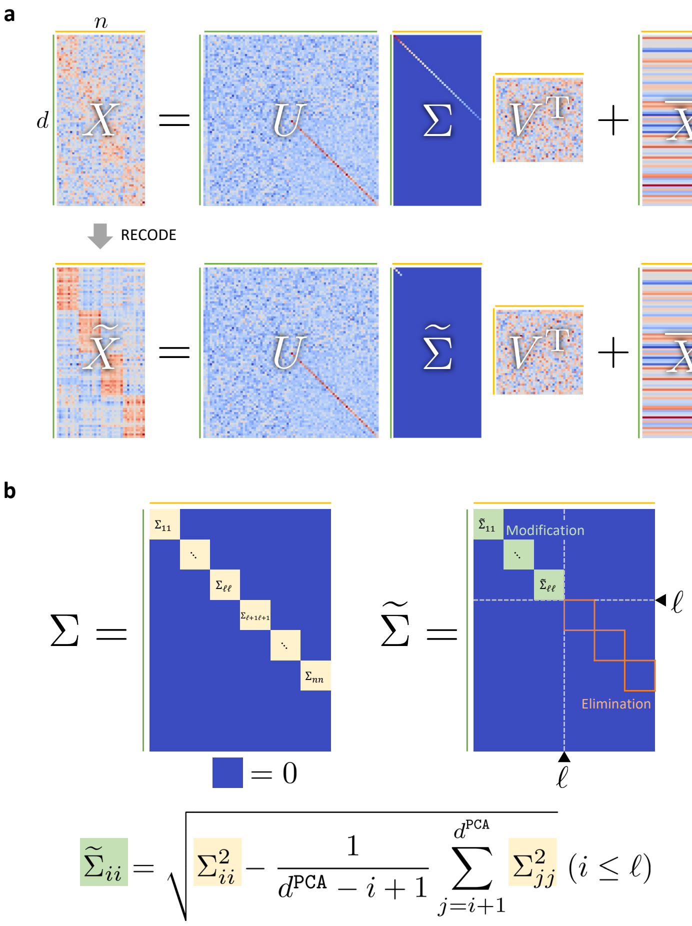


Figure ST4. Interpretation of RECODE from singular value decomposition. **a**, Illustration of the decompositions of observed data matrix $X$ and RECODE-modified data matrix $\widetilde{X}$. These decompositions are derived from the singular value decompositions of the difference data, that is, $X - \overline{X} = U_X \Sigma V_X^{\mathrm{T}}$ and $\widetilde{X}_\ell - \overline{X} = U_X \widetilde{\Sigma} V_X^{\mathrm{T}}$. **b**, Relationship between the rectangular diagonal matrices $\Sigma$ and $\widetilde{\Sigma}$ when the eigenvalue modification is given by the NRM.

From Theorem 3.5, because the second term on the right-hand side of Eq. (13) does not depend on the dimension $d$, the transformation (14) prevents the divergence of the distances between the observed data. This is in contrast to the loss of true information shown in Proposition 1.1. Consequently, we can expect a smaller error in the observed distance for a smaller $m$. That is, the minimum $m$ for which condition (C1) holds is the optimal value for the parameter $\ell$ in RECODE. With such a setting, RECODE can resolve COD1 and recover accurate measurements.

To estimate the minimum $m$ for which condition (C1) holds, we present the following theorem. The proof is provided in Appendix B.

**Theorem 3.6.** *For $m$ satisfying condition (C1),*

$$\lambda_{X,i} = u_{X,i}^{\mathrm{T}} S_E u_{X,i}$$

*holds for $i = m+1, \ldots, d$, where $S_E$ is the covariance matrix of the noise matrix $E = (e_1, \ldots, e_n)$.*

From Theorem 3.6, if the covariance matrix $S_E$ of the noise is known, we can numerically estimate the optimal value $\ell^{\mathtt{opt}}$ of the parameter $\ell$ using

$$\ell^{\mathtt{opt}} = \min\left\{ k \in \{1, \ldots, d\};\ \sum_{i=k+1}^{d} \lambda_{X,i} \le \sum_{i=k+1}^{d} u_{X,i}^{\mathrm{T}} S_E u_{X,i} \right\}, \tag{15}$$

which is an approximation of the minimum $m$ for which condition (C1) empirically. However, it is difficult to accurately estimate the covariance matrix $S_E$ of the noise from the observed data. Assuming the independence of $e_{i*}$ for $i = 1, \ldots, d$, where $e_{i*}$ is the distribution followed by $e_{i1}, \ldots, e_{in}$, the covariance matrix $S_E$ can be evaluated as $S_E = \mathrm{diag}(\mathrm{Var}(e_{1*}), \ \ldots, \ \mathrm{Var}(e_{d*}))$. Therefore, under the aforementioned assumption, if the noise variances $\mathrm{Var}(e_{i*})$ $(i = 1, \ldots, d)$ are known, then we can evaluate the optimal parameter $\ell^{\mathtt{opt}}$ using Eq. (15). In particular, if there exists $s^2 > 0$ such that $\mathrm{Var}(e_{i*}) = s^2$ for $i = 1, \ldots, d$, that is, all the noise variances are equivalent, then we have

$$\begin{aligned}
\sum_{i=k+1}^{d} u_{X,i}^{\mathrm{T}} S_E u_{X,i} &= \sum_{i=k+1}^{d} \sum_{j=1}^{d} \mathrm{Var}(e_{j*}) u_{X,ji}^2 \\
&= s^2 \sum_{i=k+1}^{d} \sum_{j=1}^{d} u_{X,ji}^2 \\
&= (d-k)s^2.
\end{aligned}$$

Consequently, under the aforementioned conditions, we can estimate the optimal value $\ell^{\mathtt{opt}}$ using

$$\ell^{\mathtt{opt}} = \min\left\{ k \in \{1, \ldots, d\};\ \sum_{i=k+1}^{d} \lambda_{X,i} \le (d-k)s^2 \right\}. \tag{16}$$

The previous argument relies on the knowledge of noise variances. Thus, we introduce the following method to estimate noise variances.

*Noise variance estimation.* We assume the following additional conditions:

(C2) The noise variances for features are equivalent, that is,

$$\exists s^2 > 0 \text{ such that } \mathrm{Var}(e_{i*}) = s^2 \quad \text{for } i = 1, \ldots, d.$$

(C3) In true data, there are some features without any variation, that is,

$$\exists k\ (1 \ll k \le d), \exists I_k = \{i_1, \ldots, i_k\} \subseteq \{1, \ldots, d\} \text{ s.t. } \mathrm{Var}(x_{i*}^{\mathtt{true}}) = 0 \text{ for } i \in I_k.$$

Here, $x_{i*}^{\mathtt{true}}$ is the distribution followed by $x_{i1}^{\mathtt{true}}, \ldots, x_{in}^{\mathtt{true}}$.

From conditions (C2)–(C3), we can evaluate the noise variance $s^2$ as

$$s^2 \approx s_{X^\star,i}^2 \quad \text{for } i \in I_k,$$

where $s^2_{X^\star,i}$ are the sample variances and the symbol $\star$ indicates the sample values. To verify this, we note that if $i \in I_k$, then $\mathrm{Var}(x^{\mathtt{true}}_{i*}) = 0$, and $s^2_{X^\star,i} \approx \mathrm{Var}(x_{i*}) = \mathrm{Var}(e_{i*}) = s^2$ consequently.

Therefore, if $k$ is sufficiently large, the mode of the sample variances becomes $s^2$. This mode can be estimated by constructing a histogram of the variances, that is,

$$
\begin{aligned}
(17) \qquad \widetilde{s}^2 &= \frac{\widetilde{k}\Delta^{\mathtt{bin}} + (\widetilde{k}+1)\Delta^{\mathtt{bin}}}{2}, \\
\widetilde{k} &:= \operatorname*{arg\,max}_{k \in \mathbb{N}} \# \left\{ i = 1, \ldots, d \;\middle|\; k\Delta^{\mathtt{bin}} \leq s^2_{X^\star,i} < (k+1)\Delta^{\mathtt{bin}} \right\}.
\end{aligned}
$$

Here, $\mathbb{N}$ is the set of nonnegative integers, and $\Delta^{\mathtt{bin}}$ is the bin size for the histogram.

For example, Fig ST5 shows a scatter plot and a histogram of the variances of the test data in Section 3.4 with a sample size of 1,000 and dimension 20,000. The dashed lines indicate the estimated noise variance $\widetilde{s}^2$. Here, the bin size $\Delta^{\mathtt{bin}}$ for the histogram is automatically set using the histogram binwidth optimization method [16].

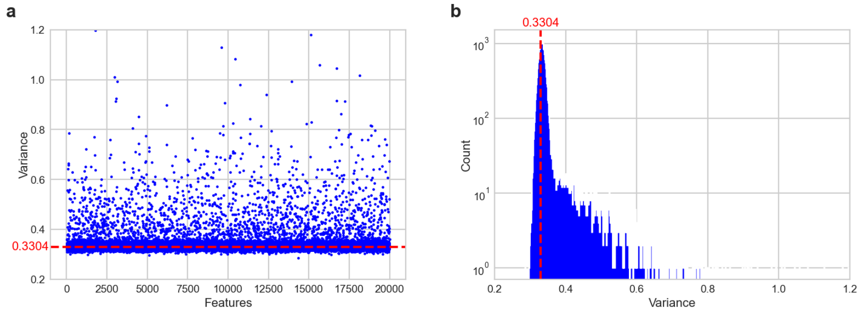


Figure ST5. Variance plots of test data in Section 3.4. **a**, Scatter plot of variances and features. **b**, Histogram of variances. The red dashed lines indicate the estimated value $\widetilde{s}^2$ of the noise variance. The true noise variance $s^2$ is 1/3.

3.3. **Computation.** In Fig ST6 and Fig ST7, we show the algorithms of RECODE and RECODE with parameter estimation, respectively. The procedures (R1)–(R3) and (E1)–(E2) are given as follows:

(R1) Compute the PCA projection $U^{\mathrm{T}}_{X^\star}(\,\cdot - \overline{X^\star})$ for the sample data matrix $X^\star$ by solving the eigenvalue problem $S_{X^\star} u_{X^\star,i} = \lambda_{X^\star,i} u_{X^\star,i}$ $(i = 1, \ldots, d)$, in which the eigenvalues satisfy $\lambda_{X^\star,1} \geq \cdots \geq \lambda_{X^\star,d}$ (e.g., by using singular value decomposition methods).

(R2) Compute the diagonal matrix $\widetilde{\Lambda}^{1/2}_{X^\star,\ell} = \mathrm{diag}(\widetilde{\lambda}_{X^\star,1}, \ldots, \widetilde{\lambda}_{X^\star,\ell}, 0, \ldots, 0)$, where modified eigenvalues $\widetilde{\lambda}_{X^\star,i}$ $(i = 1, \ldots, \ell)$ are computed by the NRM (4).

(R3) Compute the RECODE-modified data matrix $\widetilde{X}^\star$ using (11), that is,

$$\widetilde{X}^\star = U_{X^\star} \widetilde{\Lambda}^{1/2}_{X^\star,\ell} \Lambda^{-1/2}_{X^\star} U^{\mathrm{T}}_{X^\star}(X^\star - \overline{X^\star}) + \overline{X^\star}.$$

(E1) Estimate the noise variance $\widetilde{s}^2$ using the noise variance estimation (17).

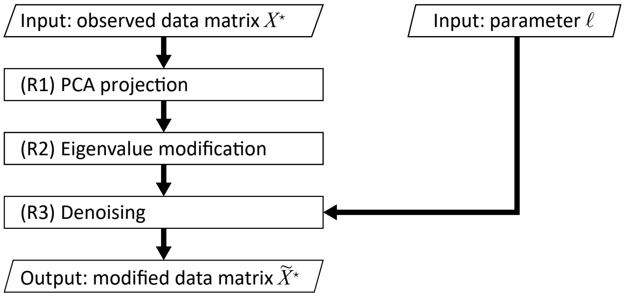


Figure ST6. Flowchart of RECODE.

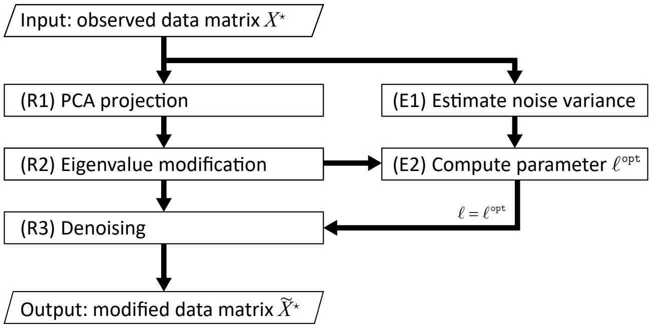


Figure ST7. Flowchart of RECODE with parameter estimation.

(E2) Compute the optimal parameter $\ell^{\mathtt{opt}}$ using (16).

Recall that the symbol $\star$ indicates sample values.

3.4. **Verification.** We investigate the COD and verify its resolution via RECODE using PCA, multidimensional scaling (MDS), t-distributed stochastic neighbor embedding (t-SNE), and uniform manifold approximation and projection (UMAP), which are standard data analysis methods for scRNA-seq [12, 13, 17].

The test data are set as follows. We sample 1,000 points from two figure-eight shapes in three-dimensional space as reference data (Fig ST8). The reference data are embedded in 20,000-dimensional space to satisfy condition (C3) with $k = 18{,}000$, yielding the true data. The observed data are set as the true data with noise following a uniform distribution in $[-1, 1]$ for each feature. Note that the rank of the true data is 3, and the noise variance $\mathrm{Var}(e_{i*})$ is 1/3 for $i = 1, \ldots, d$. Moreover, in this setting, noise does not affect the PCs in the PCA. Therefore, conditions (C1)–(C3) are satisfied.

We apply RECODE with parameter estimation, as shown in Fig ST7, to the observed data. In the process of noise variance estimation (E1), the estimated

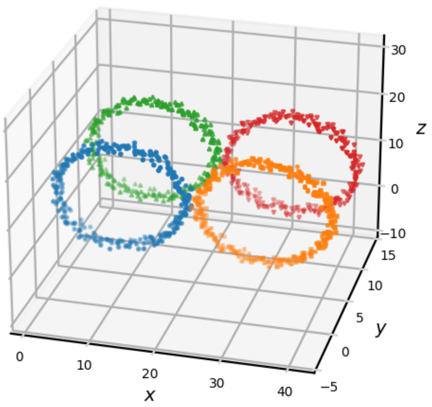


Figure ST8. Reference data (two figure-eight shapes).

value $\widetilde{s}^2$ of the noise variance is evaluated as 0.3304 (see Fig ST5). In the process of parameter estimation (E2), the optimal parameter $\ell^{\mathtt{opt}}$ is evaluated as 8 (Fig ST9).

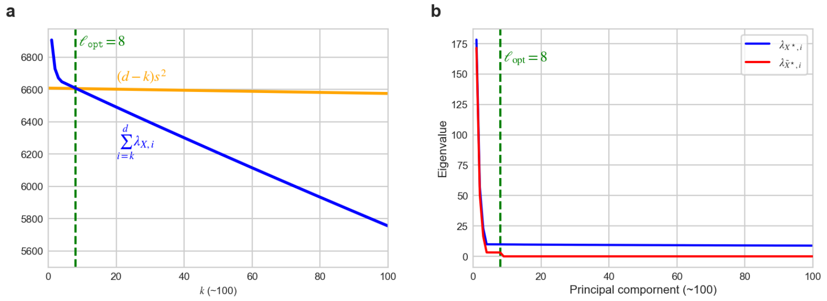


Figure ST9. Parameter estimation and modified eigenvalue. **a**, Estimation of optimal parameter $\ell^{\mathtt{opt}}$ using Eq. (16). **b**, Modified eigenvalues $\lambda_{\widetilde{X},i}$ $(1 \leq i \leq 100)$ by RECODE.

Fig ST10 shows the two-dimensional projections by PCA, MDS, t-SNE, and UMAP for the true data, observed data, and RECODE-modified data. As the computations for MDS, t-SNE, and UMAP involve the computation of distances, COD1 (loss of closeness) renders them unable to recover the correct structure of the reference data. We also consider the cumulative contribution rates (Fig ST11 and Table ST1) of the PCs of the PCA. As explained in Section 1.2, the contribution rates of PCA are affected by COD2 (inconsistency of statistics). However, the correct structure of the data and contribution rates are recovered using RECODE (Fig ST11 and Table ST1). Consequently, we conclude that RECODE can resolve COD1 and COD2.

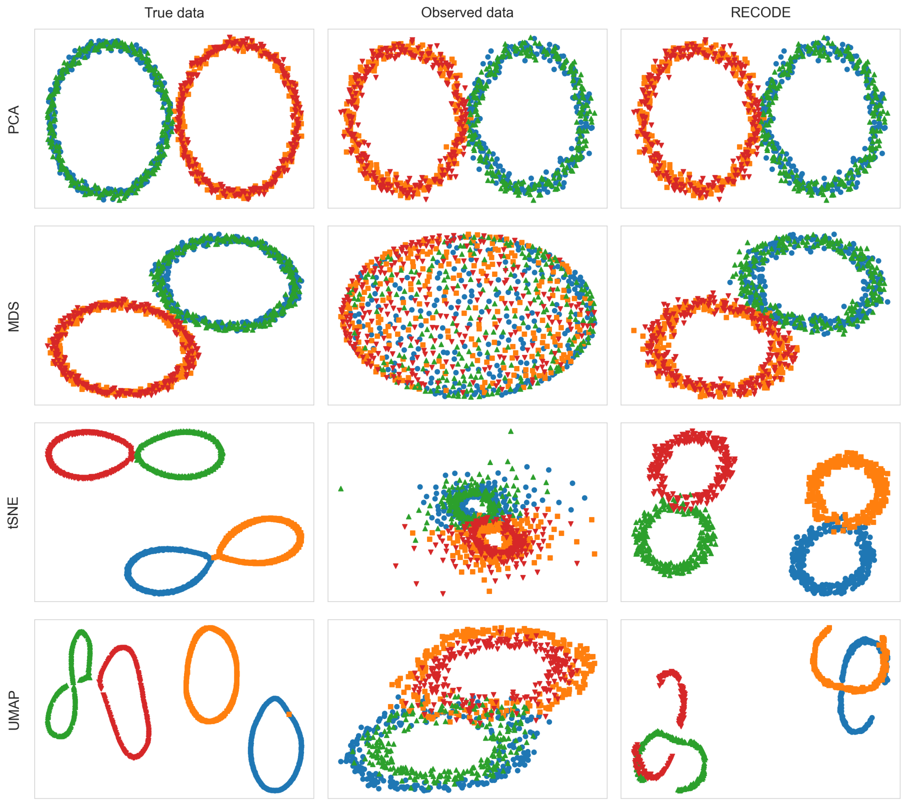


Figure ST10. Two-dimensional projections by PCA, MDS, t-SNE, and UMAP for true data, observed data, and RECODE-modified data. Colors correspond to those used in the reference data in Fig ST8.

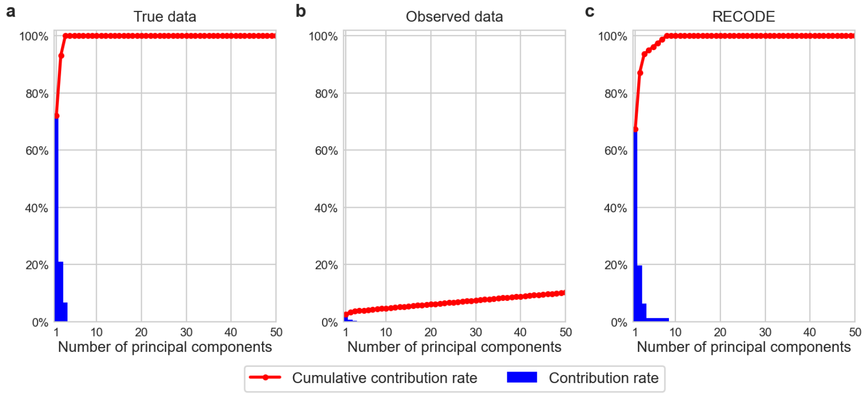


Figure ST11. Contribution rates (blue) and cumulative contribution rates (red) for the first 50 PCs of PCA.

Table ST1. Contribution rates of PCA.

| | PC1 | PC2 | PC3 |
|---|---|---|---|
| True data | 72.11% | 21.07% | 6.82% |
| Observed data | 2.58% | 0.83% | 0.34% |
| RECODE | 67.40% | 19.78% | 6.47% |

#### 4. RECODE for single-cell RNA-sequencing data

In this section, we propose an extension of RECODE to scRNA-seq data represented by unique molecular identifier (UMI) counts. Specifically, scRNA-seq involves the examination of the entire RNA information of single cells. As scRNA-seq technology reacts to and sequences randomly selected RNA molecules in single cells, scRNA-seq data involve noise from random sampling. Generally, the variance of noise from random sampling increases in proportion to the true expression values. Therefore, the variations in genes with low expression values are hidden by those in genes with high expression values. Moreover, scRNA-seq data do not satisfy the condition that the variances of noise are constant; meeting such a condition is required in parameter estimation in RECODE, as explained in Section 3.2. To resolve this issue, we analyze the statistics of noise in scRNA-seq data and propose an extension of RECODE that involves pre- and postprocessing.

4.1. **Statistical theory and method.** We introduce the notation used in this subsection. For each pair $(i, j)$ $(i = 1, \ldots, d, j = 1, \ldots, n)$, let $c_{ij}$ and $c_{ij}^{\mathtt{true}}$ be random variables representing the UMI count and RNA count of gene $i$ in cell $j$, respectively. Note that $c_{ij}$ and $c_{ij}^{\mathtt{true}}$ are nonnegative integers. Let $t_j$ and $t_j^{\mathtt{true}}$ be the total counts of UMI and RNA of cell $j$; that is,

$$t_j = \sum_{i=1}^{d} c_{ij} \quad \text{and} \quad t_j^{\mathtt{true}} = \sum_{i=1}^{d} c_{ij}^{\mathtt{true}},$$

respectively. As the scales of $c_{ij}$ and $c_{ij}^{\mathtt{true}}$ can vary significantly, we consider the observed and true data scaled as $x_{ij} = c_{ij}/t_j$ and $x_{ij}^{\mathtt{true}} = c_{ij}^{\mathtt{true}}/t_j^{\mathtt{true}}$, respectively. Then, the noise $e_{ij}$ of gene $i$ for cell $j$ is defined as follows:

$$e_{ij} = x_{ij} - x_{ij}^{\mathtt{true}}.$$

Next, we consider the distribution of the UMI count data. The procedure for obtaining scRNA-seq data consists of six technical steps (Fig ST12), which can be divided into three types: copying, amplification, and sequencing. The copying and sequencing steps consist of random sampling errors because they involve randomly selecting molecules. In contrast, the amplification step involves an error caused by amplification, which is different for each molecule. UMI count data are known to be free from amplification errors because of the use of UMI tags [10]. In the UMI count data, each RNA is given a different UMI tag before amplification. This cancels the amplification errors. Therefore, we assume that the UMI count data are mainly affected by random sampling errors.

In this subsection, we recall that the random variables $c_{ij}$, which represent the UMI counts, are i.i.d. for $j = 1, \ldots, n$. When $t_j$ and $x_{ij}^{\mathtt{true}}$ are given, the probability of $c_{ij} = k$ can be regarded as the probability of obtaining exactly $k$ successes in

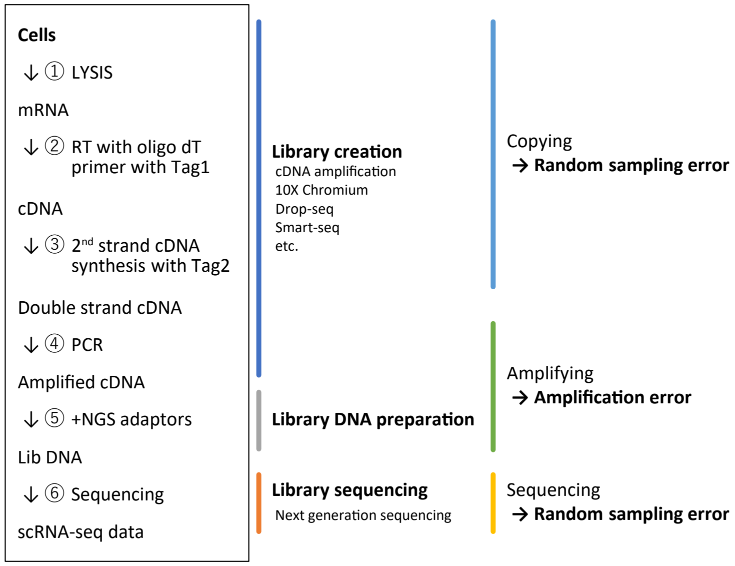


Figure ST12. Workflow of scRNA-seq and classifications of data sampling processes.

$t_j$ independent Bernoulli trials with probability $x_{ij}^{\texttt{true}}$. Then, the observed data $c_{ij}$ follow a binomial distribution conditioned on the values of $t_j$ and $x_{ij}^{\texttt{true}}$:

$$\Pr(c_{ij} = k | t_j = t, x_{ij}^{\texttt{true}} = y) = {}_t\mathrm{C}_k y^k (1-y)^{t-k}, \quad k = 0, 1, \ldots,$$

where ${}_a\mathrm{C}_b$ is the binomial coefficient

$${}_a\mathrm{C}_b = \binom{a}{b} = \frac{a!}{(a-b)!b!}$$

indexed by $a \geq b \geq 0$. Here, because the dimension $d$ is large, each RNA count $c_{ij}^{\texttt{true}}$ is small relative to the total number of RNA counts $t_j^{\texttt{true}}$. It is reasonable to assume that the ratio $x_{ij}^{\texttt{true}} = c_{ij}^{\texttt{true}} / t_j^{\texttt{true}}$ of the RNA content is small. Accordingly, under the assumption that the total number of UMI counts $t_j$ is sufficiently large, the UMI counts $c_{ij}$ approximately follow a Poisson distribution with parameter $t_j x_{ij}^{\texttt{true}}$. Thus, in the following, the conditional probability of $c_{ij}$ is assumed to be a Poisson distribution:

$$(18) \qquad \Pr(c_{ij} = k | t_j = t, x_{ij}^{\texttt{true}} = y) = \mathrm{Poisson}(k; \lambda = ty) = \frac{(ty)^k e^{-ty}}{k!}.$$

As discussed in many studies [5, 7, 10, 11, 18], UMI counts $c_{ij}$ are known to be related to Poisson distributions. However, we note that the parameter $\lambda$ depends on the random variables $t_j$ and $x_{ij}^{\texttt{true}}$; hence, $c_{ij}$ do not necessarily follow a Poisson distribution. In fact, we may observe larger variances than those when a Poisson distribution is assumed. These phenomena are often called overdispersions. To explain these phenomena, distributions such as a gamma or negative binomial distribution have been considered [5, 7]. However, these distributions require the fitting of parameters in order to infer statistics.

In contrast, using our analysis, we can derive the variance $\mathrm{Var}(x_{ij})$ of scaled data without using any parameters, as follows:

**Theorem 4.1.** *Under the conditional probability* (18),

$$\mathrm{Var}(x_{ij}) = \mathrm{Var}(x_{ij}^{\texttt{true}}) + \mathrm{E}(x_{ij}/t_j), \quad i = 1, \ldots, d, \ j = 1, \ldots, n$$

*holds.*

We provide a proof in Appendix B. A previous study [7] and its related literature claimed that the noise variance $\mathrm{Var}(e_{ij})$ is explained by the mean $\mathrm{E}(x_{ij})$ of the scaled data $x_{ij} = c_{ij}/t_j$. In contrast, we emphasize that when $x_{ij}^{\texttt{true}}$ is constant, the noise variance can be expressed as $\mathrm{Var}(e_{ij}) = \mathrm{E}(x_{ij}/t_j)$ from Theorem 4.1. That is, the noise variance $\mathrm{Var}(e_{ij})$ of the scaled data is explained by the mean $\mathrm{E}(x_{ij}/t_j)$ of the square-scaled data $x_{ij}/t_j = c_{ij}/t_j^2$. Therefore, we should examine the $\mathrm{E}(x_{ij}/t_j)$–$\mathrm{Var}(x_{ij})$ relationship rather than the mean–variance relationship, which is verified in Section 4.3 (see also Fig ST15).

We now introduce the following classification of genes:

$$\begin{cases} i\text{th gene is } \textit{significant} & \overset{\texttt{def}}{\Longleftrightarrow} x_{ij}^{\texttt{true}} \neq \text{const. for } j = 1, \ldots, n, \\ i\text{th gene is } \textit{non-significant} & \overset{\texttt{def}}{\Longleftrightarrow} x_{ij}^{\texttt{true}} = \text{positive const. for } j = 1, \ldots, n, \\ i\text{th gene is } \textit{silent} & \overset{\texttt{def}}{\Longleftrightarrow} x_{ij}^{\texttt{true}} = 0 \text{ for } j = 1, \ldots, n. \end{cases}$$

From this definition, significant genes capture cell-identifiable features, whereas the non-significant genes (e.g., housekeeping genes) do not identify cell differences. Silent genes are those that have no functions.

To infer the classification of each gene based on its distribution, we show the relationships between their means and variances. Let $I^{\texttt{sig}}$, $I^{\texttt{non-sig}}$, and $I^{\texttt{silent}}$ be the index sets of significant, non-significant, and silent genes, respectively. From the definition of gene classification, for $j = 1, \ldots, n$, we have

$$\mathrm{E}(x_{ij}^{\texttt{true}}) \begin{cases} > 0, & i \in I^{\texttt{sig}}, \\ > 0, & i \in I^{\texttt{non-sig}}, \\ = 0, & i \in I^{\texttt{silent}}, \end{cases} \quad \mathrm{Var}(x_{ij}^{\texttt{true}}) \begin{cases} > 0, & i \in I^{\texttt{sig}}, \\ = 0, & i \in I^{\texttt{non-sig}}, \\ = 0, & i \in I^{\texttt{silent}}. \end{cases}$$

Therefore, from Theorem 4.1, for $j = 1, \ldots, n$, we obtain the following corollary:

**Corollary 4.2.** *Under the condition of Theorem 4.1, for* $j = 1, \ldots, n$,

$$\mathrm{Var}(x_{ij}) = \begin{cases} \mathrm{Var}(x_{ij}^{\texttt{true}}) + \mathrm{E}(x_{ij}/t_j), & i \in I^{\texttt{sig}}, \\ \mathrm{E}(x_{ij}/t_j), & i \in I^{\texttt{non-sig}}, \\ 0, & i \in I^{\texttt{silent}} \end{cases}$$

*holds.*

As stated earlier, non-significant genes cannot be used to identify differences among target cells. This implies that their distribution is a translation of the noise distribution. Thus, from Corollary 4.2, for $i \in I^{\texttt{non-sig}}$, the noise variances are explained using the means of $x_{ij}/t_j$.

Furthermore, we note that the variances of non-significant genes with high expression values may be much larger than those of significant genes with low expression values, that is, for $i \in I^{\texttt{sig}}$ and $i' \in I^{\texttt{non-sig}}$,

$$\mathrm{Var}(x_{ij}) = \mathrm{Var}(x_{ij}^{\texttt{true}}) + \mathrm{E}(x_{ij}/t_j) \ll \mathrm{E}(x_{i'j}/t_j) = \mathrm{Var}(x_{i'j}). \tag{19}$$

This causes a problem in PCA; that is, condition (C1) may not be satisfied. In particular, the PCA may extract the $i'$th non-significant gene as a principal component. We call this third problem of the COD in scRNA-seq data the *inconsistency of principal components* (COD3).

On the basis of the aforementioned discussions, we present a normalization to make the noise variances constant. Let $f_\alpha : \mathbb{R} \to \mathbb{R}$ be a function

$$f_\alpha(x) := \begin{cases} \dfrac{x}{\sqrt{\alpha}}, & \alpha > 0, \\ 0, & \alpha = 0 \end{cases}$$

parameterized by $\alpha \geq 0$, with inverse

$$f_\alpha^{-1}(x) = \sqrt{\alpha}x.$$

Then, we define a normalized random variable $z_{ij}$ as

$$z_{ij} := f_{\mathrm{E}(x_{ij}/t_j)}(x_{ij}).$$

We call this normalization a *noise variance-stabilizing normalization* (NVSN), the terminology of which is justified below. From Corollary 4.2, it follows that the variance of $z_{ij}$ is

$$\mathrm{Var}(z_{ij}) = \begin{cases} \mathrm{Var}(z_{ij}^{\texttt{true}}) + 1, & i \in I^{\texttt{sig}}, \\ 1, & i \in I^{\texttt{non-sig}}, \\ 0, & i \in I^{\texttt{silent}}, \end{cases} \tag{20}$$

where $z_{ij}^{\texttt{true}} := f_{\mathrm{E}(x_{ij}/t_j)}(x_{ij}^{\texttt{true}})$. Thus, the following inequality holds for significant and non-significant genes:

$$\mathrm{Var}(z_{ij}) \geq 1, \qquad i \in I^{\texttt{sig}} \cup I^{\texttt{non-sig}}. \tag{21}$$

Let the noise of $z_{ij}$ be denoted by

$$e'_{ij} = z_{ij} - z_{ij}^{\texttt{true}}.$$

In scRNA-seq data creation, it is considered that the noise derived from random sampling occurs independently from the behavior of true data, meaning that $e'_{ij}$ and $z_{ij}^{\texttt{true}}$ may be independent. Then, from $\mathrm{Var}(z_{ij}) = \mathrm{Var}(z_{ij}^{\texttt{true}}) + \mathrm{Var}(e'_{ij})$, we have

$$\mathrm{Var}(e'_{ij}) = \begin{cases} 1, & i \in I^{\texttt{sig}} \cup I^{\texttt{non-sig}}, \\ 0, & i \in I^{\texttt{silent}}. \end{cases} \tag{22}$$

Therefore, NVSN stabilizes the noise variances such that they are constant. Note that NVSN is different from the well-known variance-stabilizing transformation (z-score transformation), which makes the variances of *all* features constant.

The baseline of the variances $\mathrm{Var}(z_{ij})$ for $I^{\texttt{sig}} \cup I^{\texttt{non-sig}}$ is set to one, and the variances only in $I^{\texttt{sig}}$ take values larger than one. This ensures that the inequality (19) never occur, that is, for all $i \in I^{\texttt{sig}}$ and $i' \in I^{\texttt{non-sig}}$,

$$\mathrm{Var}(z_{ij}) = \mathrm{Var}(z_{ij}^{\texttt{true}}) + 1 > 1 = \mathrm{Var}(z_{i'j}).$$

Thus, the normalized data matrix $Z = (z_{ij})$ are expected to satisfy condition (C1) with a sufficiently small $m$. Therefore, the normalized data matrix $Z = (z_{ij})$ are more suitable for regular RECODE than the original UMI count data matrix $C = (c_{ij})$ or the scaled data matrix $X = (x_{ij})$. Moreover, because the noise variance is explicitly given by Eq. (22), we can use parameter optimization (15) without the noise variance estimation. Furthermore, by eliminating the silent genes in advance, we can use the simple form (16) of parameter optimization because all the noise variances of the genes are equal to one ($s^2 = 1$).

After we derive the modified data matrix $\widetilde{Z} = (\widetilde{z}_{ij})$ by applying regular RECODE (11) to the normalized data matrix $Z = (z_{ij})$, the modified UMI count data matrix $\widetilde{C} = (\widetilde{c}_{ij})$ are obtained by rescaling

$$\widetilde{x}_{ij} = f^{-1}_{\mathrm{E}(x_{ij}/t_j)}(\widetilde{z}_{ij}),$$
$$\widetilde{c}_{ij} = t_j \widetilde{x}_{ij}.$$

We summarize the formulation of RECODE for scRNA-seq data as follows.

**RECODE for scRNA-seq data**

For the scRNA-seq data matrix $C = (c_{ij}) \in \mathbb{R}^{d\times n}$ without silent genes, RECODE defines the modified data matrix

$$\widetilde{C} \coloneqq F^{-1}(U_{F(C)}\widetilde{\Lambda}^{1/2}_{F(C),\ell^{\mathrm{opt}}}\Lambda^{-1/2}_{F(C)}U^{\mathrm{T}}_{F(C)}[F(C) - \overline{F(C)}] + \overline{F(C)}).$$

Here,

$$F : \mathbb{R}^{d\times n} \to \mathbb{R}^{d\times n} \text{ such that } [F(C)]_{ij} \coloneqq f_{\mathrm{E}(c_{ij}/t_j^2)}(c_{ij}/t_j),$$

$$\ell^{\mathtt{opt}} \coloneqq \min\left\{k \in \{1,\dots,d\};\ \sum_{i=k+1}^{d} \lambda_{F(C),i} \le (d-k)\right\}.$$

4.2. **Computation.** We now explain the computations of RECODE to sample scRNA-seq data. We use the symbol $\star$ to indicate sample values; for example, $C^\star = (c^\star_{ij})$ $(i = 1,\dots,d,\ j = 1,\dots,n)$ for the sample UMI count data matrix. We remove silent genes such that $c^\star_{ij} = 0$ for $j = 1,\dots,n$ in advance from the sample UMI count data, that is, $\{1,\dots,d\} = I^{\mathtt{sig}} \cup I^{\mathtt{non\text{-}sig}}$ and $I^{\mathtt{silent}} = \emptyset$. The total UMI counts $t^\star_j$ and scaled data matrix $X^\star = (x^\star_{ij})$ $(i = 1,\dots,d,\ j = 1,\dots,n)$ are computed as

$$t^\star_j = \sum_{i=1}^{d} c^\star_{ij} \quad \text{and} \quad x^\star_{ij} = \frac{c^\star_{ij}}{t^\star_j},$$

respectively. We define $\mu_{(X/T)^\star,i}$ as

$$\mu_{(X/T)^\star,i} \coloneqq \frac{1}{n}\sum_{j=1}^{n}\frac{x^\star_{ij}}{t^\star_j}.$$

Note that $\mu_{(X/T)^\star,i}$ is the sample mean of square-scaled data $x^\star_{ij}/t^\star_j = c^\star_{ij}/(t^\star_j)^2$, which corresponds to an estimate of $\mathrm{E}(x_{ij}/t_j)$ and represents the estimated noise variance. Then, the normalized data matrix $Z^\star = (z^\star_{ij})$ is computed as

$$z^\star_{ij} = f_{\mu_{(X/T)^\star,i}}(x^\star_{ij}), \quad i = 1,\dots,d,\ j = 1,\dots,n. \tag{23}$$

We apply regular RECODE (11) to the normalized data matrix $Z^\star = (z^\star_{ij})$; that is, we compute the RECODE-modified normalized data matrix

$$\widetilde{Z}^\star = U_{Z^\star}\widetilde{\Lambda}^{1/2}_{Z^\star,\ell^{\mathrm{opt}}}\Lambda^{-1/2}_{Z^\star}U^{\mathrm{T}}_{Z^\star}(Z^\star - \overline{Z^\star}) + \overline{Z^\star}. \tag{24}$$

Here, because the silent genes have already been removed, all noise variances can be estimated as $\mathrm{Var}(e_{i*}) = 1$ $(i = 1, \ldots, d)$. Therefore, the covariance matrix $S_E$ in the evaluation of the optimal parameter (15) is evaluated as the identity matrix, that is,

$$S_E = I_d.$$

Consequently, we can compute the optimal parameter $\ell^{\mathtt{opt}}$ using

$$\ell^{\mathtt{opt}} = \min\left\{ k \in \{1, \ldots, d\};\ \sum_{i=k+1}^{d} \lambda_{Z^\star,i} \leq (d-k) \right\}. \tag{25}$$

After applying RECODE, the modified data matrix $\widetilde{X}^\star = (\widetilde{x}^\star_{ij})$ and modified UMI count data matrix (RECODE-modified data matrix) $\widetilde{C}^\star = (\widetilde{c}^\star_{ij})$ are evaluated as follows:

$$\widetilde{x}^\star_{ij} = f^{-1}_{s^2_{E,i}}(\widetilde{z}^\star_{ij}), \tag{26}$$

$$\widetilde{c}^\star_{ij} = t^\star_j \widetilde{x}^\star_{ij}. \tag{27}$$

We summarize the algorithm of RECODE for scRNA-seq data as follows:

I. Compute the normalized data matrix $Z^\star = (z^\star_{ij})$ from the UMI count data matrix $C^\star = (c^\star_{ij})$ using the noise variance-stabilizing normalization (NVSN) (23).

II. Compute the PCA projection $U^{\mathrm{T}}_{Z^\star}(\cdot - \overline{Z^\star})$ for the sample data matrix $Z^\star$ by solving the eigenvalue problem $S_{Z^\star} u_{Z^\star,i} = \lambda_{Z^\star,i} u_{Z^\star,i}$ $(i = 1, \ldots, d)$, in which the eigenvalues satisfy $\lambda_{Z^\star,1} \geq \cdots \geq \lambda_{Z^\star,d}$ (e.g., by using singular value decomposition methods).

III. Compute the diagonal matrix $\widetilde{\Lambda}^{1/2}_{Z^\star,\ell^{\mathtt{opt}}} = \mathrm{diag}(\widetilde{\lambda}_{Z^\star,1}, \ldots, \widetilde{\lambda}_{Z^\star,\ell^{\mathtt{opt}}}, 0, \ldots, 0)$, where modified eigenvalues $\widetilde{\lambda}_{Z^\star,i}$ $(i = 1, \ldots, \ell^{\mathtt{opt}})$ and optimal parameter $\ell^{\mathtt{opt}}$ are computed by the NRM (4) and Eq. (25), respectively.

IV. Compute the RECODE-modified normalized data matrix $\widetilde{Z}^\star$ by Eq. (24). Furthermore, compute the RECODE-modified data matrix $\widetilde{C}^\star = (\widetilde{c}^\star_{ij})$ using Eqs. (26) and (27). If there exist negative values in $\widetilde{C}^\star = (\widetilde{c}^\star_{ij})$, these negative values are modified to zero.

The flow chart is shown in Fig ST13 (see also Fig 3a in the main manuscript).

4.3. **Verification by scRNA-seq data.** In this section, we test RECODE numerically using the "3k PBMCs from a Healthy Donor" data, an example of scRNA-seq data represented by UMI counts; these data are obtained from the single-cell gene expression datasets provided by 10X Genomics, Inc. [6]. The library creation of scRNA-seq data is conducted with 10X Chromium with version 3.1 chemistry. The sample data contain 2,700 cells ($n$ = 2,700) and 32,738 genes with 16,104 silent genes. Thus, the dimensions of the data without silent genes are 16,634 ($d$ = 16,634).

We apply RECODE to the scRNA-seq data (Fig ST14). The optimal parameter value $\ell^{\mathtt{opt}}$, calculated using Eq. (25), is 439 (Fig ST14d). The numbers of significant and non-significant genes are 8,275 (49.7%) and 8,359 (50.3%), respectively. Here,

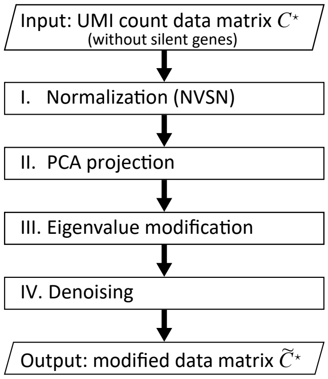


Figure ST13. Flowchart of RECODE for scRNA-seq data.

we determine the significance of the genes by the magnitude of the sample variance $s^2_{Z^\star,i}$ of the normalized data as

$$\begin{aligned} s^2_{Z^\star,i} > 1 &\Leftrightarrow \text{ significant } (i \in I^{\texttt{sig}}), \\ s^2_{Z^\star,i} \leq 1 &\Leftrightarrow \text{ non-significant } (i \in I^{\texttt{non-sig}}). \end{aligned}$$

Here, $s^2_{Y^\star,i}$ is the sample variance of feature $i$ for the sample data matrix $Y^\star = (y^\star_{ij})$, that is,

$$s^2_{Y^\star,i} = \frac{1}{n-1}\sum_{j=1}^{n}(y^\star_{ij} - \mu_{Y^\star,i})^2,$$

and $\mu_{Y^\star,i}$ is the sample average, that is,

$$\mu_{Y^\star,i} = \frac{1}{n}\sum_{j=1}^{n} y^\star_{ij}.$$

We first examine the variance model derived by Theorem 4.1. Fig ST15 shows the relationship among the sample statistics of the genes: (a) the sample mean $\mu_{X^\star,i}$ of scaled data $c^\star_{ij}/t^\star_j$ versus the sample variance $s^2_{X^\star,i}$ of scaled data and (b) the sample mean $\mu_{(X/T)^\star,i}$ of square-scaled data $c^\star_{ij}/(t^\star_j)^2$ versus the sample variance $s^2_{X^\star,i}$ of scaled data. The blue crosses denote the values of the housekeeping genes that code the ribosomal proteins and mitochondrial ribosomal proteins (224 genes), which are biologically categorized as non-significant genes. Suppose that the true value $x^{\texttt{true}}_{ij}$ and the noise $e_{ij}$ are independent and that the noise variance $\mathrm{Var}(e_{ij})$ is explained by a variable $\xi_{ij}$, that is, $\mathrm{Var}(e_{ij}) = \varphi(\xi_{ij})$ and $\mathrm{Var}(x_{ij}) = \mathrm{Var}(x^{\texttt{true}}_{ij}) + \varphi(\xi_{ij})$ ($\varphi$: a function of $\xi_{ij}$). Then, it holds that

$$\mathrm{Var}(x_{ij}) \geq \mathrm{Var}(x_{i'j}) \text{ for } i \in I^{\texttt{sig}} \cup I^{\texttt{non-sig}} \text{ and } i' \in I^{\texttt{non-sig}} \text{ s.t. } \xi_{ij} = \xi_{i'j}.$$

In other words, the variances of the non-significant genes become minimal for each value of $\xi_{ij}$. In the case of Fig ST15a ($\xi_{ij} = \mathrm{E}(x_{ij})$), which corresponds to conventional studies that consider Poisson, gamma, and negative binomial distributions, the variances of some housekeeping genes (blue crosses) are not the minimum values. In contrast, in the case of Fig ST15b ($\xi_{ij} = \mathrm{E}(x_{ij}/t_j)$), most variances of

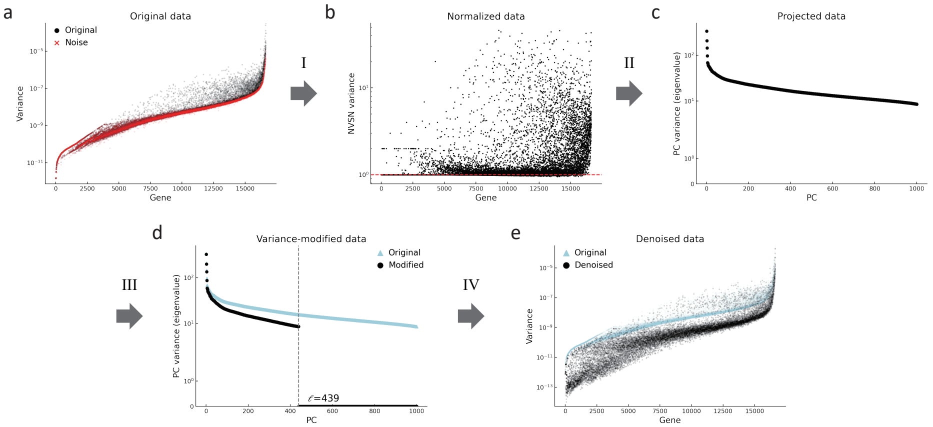


Figure ST14. Variance (eigenvalue) distributions in the computational processes in RECODE. The notations over the arrows correspond to those in the flowchart in Fig ST13. The genes on the horizontal axes (**a**, **b**, **e**) are sorted by means of scaled data. The PC on the horizontal axes (**c**, **d**) denote the principal component.

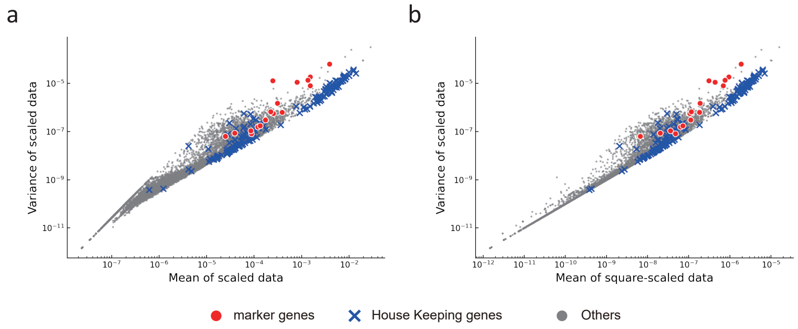


Figure ST15. Relationships among sample statistics of genes. **a**, sample mean of scaled data versus the sample variance of scaled data; **b**, sample mean of square-scaled data versus the sample variance of scaled data. The red circles and blue crosses denote the values of the marker genes of PBMC (*IL7R*, *CD79A*, *MS4A1*, *CD8A*, *CD8B*, *LYZ*, *CD14*, *LGALS3*, *S100A8*, *GNLY*, *NKG7*, *KLRB1*, *FCGR3A*, *MS4A7*, *FCER1A*, *CST3*, and *PPBP*) and the housekeeping genes that code ribosomal proteins and mitochondrial ribosomal proteins, respectively.

housekeeping genes attain the minimum values; this fact is based on the aforementioned inequality. Consequently, the mean $\mathrm{E}(x_{ij}/t_j)$ of the square-scaled data can explain the noise variances more precisely.

Next, we examine the effectiveness of NVSN. Recall that Fig ST14b shows the variances $s^2_{Z^\star,i}$ of normalized data for each gene. The normalized data matrix $Z^\star$ satisfy $s^2_{Z^\star,i} \geq 1$ $(i = 1, \ldots, d)$ for almost all genes $i$, corresponding to Eq. (21), and some of them are distributed around one. In the biological setting, we can suppose that the number of non-significant genes is relatively large, meaning that $s^2_{Z^\star,i} = 1$ is satisfied for most genes. This implies that the result may provide further evidence that our modeling of noise variance (Theorem 4.1 and Fig ST14b) is optimal. These discussions lead to the applicability of RECODE, which is introduced in Section 5.2.

Next, we verify the effectiveness of RECODE by comparing the RECODE-modified data with the original data. Hereafter, in the same manner as in this study, we use $X^{\star,\mathtt{LN}} = (x^{\star,\mathtt{LN}}_{ij})$ for the original data and $\widetilde{X}^{\star,\mathtt{LN}} = (\widetilde{x}^{\star,\mathtt{LN}}_{ij})$ for the RECODE-modified data with size and log scaling, so-called log-normalize method, given by

$$x^{\star,\mathtt{LN}}_{ij} = \log_2(10^4 \times x^\star_{ij} + 1), \qquad \widetilde{x}^{\star,\mathtt{LN}}_{ij} = \log_2(10^4 \times \widetilde{x}^\star_{ij} + 1).$$

Fig ST16 shows the variance and coefficient of variation (CV) of the original and RECODE-modified data after the size and log scaling. As can be observed, the scatter plot for the original data forms a curve at the bottom of the distribution, which is actually caused by noise. In contrast, this curve is reduced to be approximately zero after applying RECODE.

Fig ST17 shows the verification of RECODE for COD1 (loss of closeness). The dendrograms of the hierarchical clustering (Fig ST17a) clearly show that RECODE preserves large-scale (long-distance) structures and uncovers finer (short-distance) structures. From Fig ST17b, small Euclidean distances and high correlation coefficients can be observed after RECODE. Moreover, because the ranges of Euclidean distances and correlation coefficients after RECODE are wider than those of the original data, RECODE can discern finer differences among single cells.

Fig ST18 shows the verification of RECODE for COD2 (inconsistency of statistics) and COD3 (inconsistency of principal components). The contribution rates of the RECODE-modified data are higher than those of the original data owing to the modification of the eigenvalues (Fig ST18a, c. 4.52% $\to$ 14.59% in PC1; 1.60% $\to$ 5.02% in PC2; 1.15% $\to$ 3.53% in PC3). Moreover, the three major principal components are different before and after the application of RECODE; the former is correlated with the depth (total counts in each cell) that is known to be independent of biological signals (Fig ST18a, b) whereas the latter can represent the difference among the three major cell types (Fig ST18c). Furthermore, the distributions of cell-specific gene expressions become more significant (Fig ST18d). As a result, RECODE enables us to correctly infer cell classifications.

From the results in this subsection, we conclude that RECODE can appropriately resolve the curses of dimensionality in the scRNA-seq data.

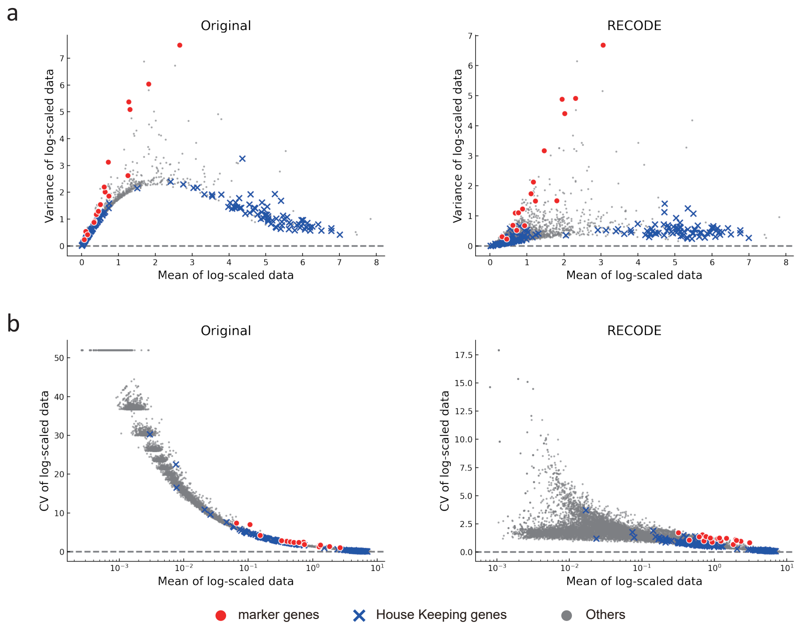


Figure ST16. Variance and coefficient of variation (CV) of the original and RECODE-modified data after size and log scaling. **a**, Scatter plots of mean versus variance. **b**, Scatter plots of mean versus CV. The red circles and blue crosses denote the values of the marker genes of PBMC and the housekeeping genes, respectively.

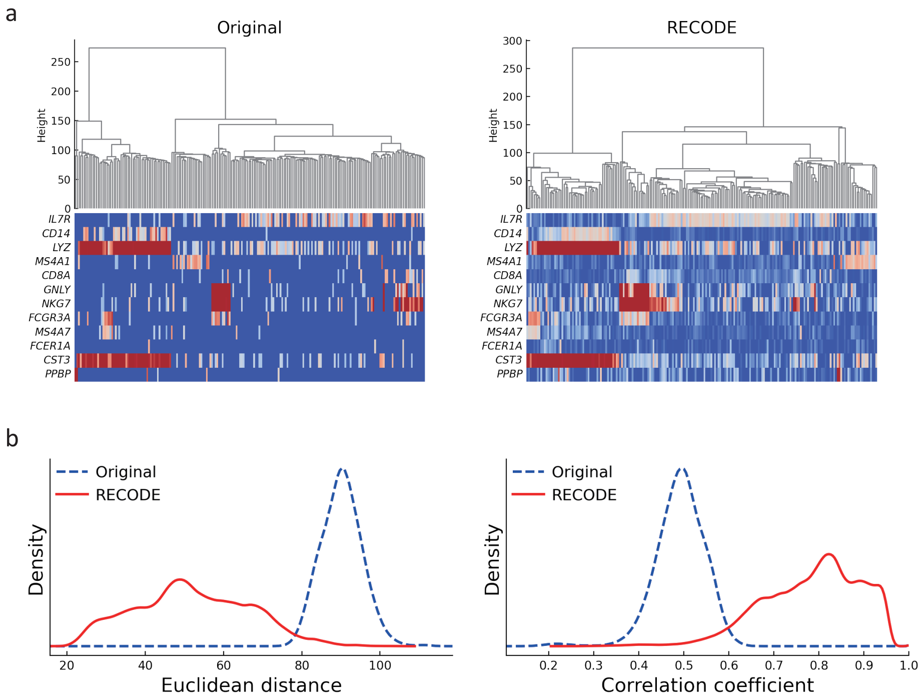


Figure ST17. Verification of RECODE for COD1 (loss of closeness). **a**, Dendrograms of hierarchical clustering based on Ward's method for 100 randomly selected cells and heat map for 12 marker genes. **b**, Density plots of the Euclidean distances and the correlation coefficients among all cells.

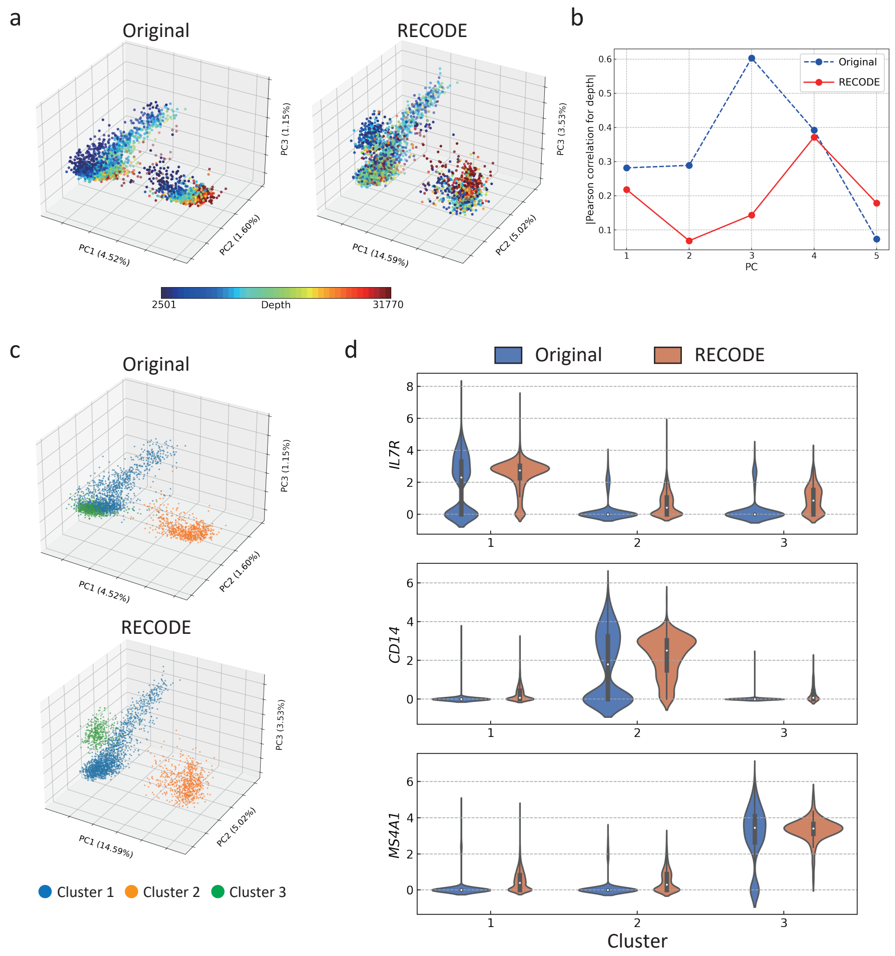


Figure ST18. Verification of RECODE for COD2 (inconsistency of statistics) and COD3 (inconsistency of principal components) **a**, PCA plots colored based on depth (total UMI counts). **b**, Absolute values of Pearson correlation for depth and principal components. **c**, PCA plots colored by clusters. **d**, Violin plot of gene expression values of marker genes (*IL7R*, *CD14*, and *MS4A1*) for clusters.

#### 5. Extensions of RECODE

5.1. **Fast algorithm.** Since $(\widetilde{\Lambda}^{1/2}_{Z^\star,\ell^{\mathrm{opt}}})_{ii} = 0$ for $i \geq \ell + 1$, we can obtain $\widetilde{Z}^\star$ by

$$\widetilde{Z}^\star = U_{Z^\star,\ell^{\mathrm{opt}}} \widehat{\Lambda}^{1/2}_{Z^\star,\ell^{\mathrm{opt}}} \Lambda^{-1/2}_{Z^\star,\ell^{\mathrm{opt}}} U^{\mathrm{T}}_{Z^\star,\ell^{\mathrm{opt}}} (Z^\star - \overline{Z^\star}) + \overline{Z^\star},$$

where $U_{Z^\star,\ell^{\mathrm{opt}}} = (u_{Z^\star,1} \dots u_{Z^\star,\ell^{\mathrm{opt}}}) \in \mathbb{R}^{d\times \ell^{\mathrm{opt}}}$, $\widehat{\Lambda}^{1/2}_{Z^\star,\ell^{\mathrm{opt}}} = \mathrm{diag}(\widetilde{\lambda}^{1/2}_{Z^\star,1}, \dots, \widetilde{\lambda}^{1/2}_{Z^\star,\ell^{\mathrm{opt}}}) \in \mathbb{R}^{\ell^{\mathrm{opt}}\times \ell^{\mathrm{opt}}}$, and $\Lambda^{-1/2}_{Z^\star,\ell^{\mathrm{opt}}} = \mathrm{diag}(\lambda^{-1/2}_{Z^\star,1}, \dots, \lambda^{-1/2}_{Z^\star,\ell^{\mathrm{opt}}}) \in \mathbb{R}^{\ell^{\mathrm{opt}}\times \ell^{\mathrm{opt}}}$. Moreover, using a matrix property

$$\sum_{i=k+1}^{d} \lambda_{Z^\star,i} = \sum_{i=1}^{d} \lambda_{Z^\star,i} - \sum_{i=1}^{k} \lambda_{Z^\star,i} = \mathrm{tr}(S_{Z^\star}) - \sum_{i=1}^{k} \lambda_{Z^\star,i},$$

we obtain

$$\ell^{\mathrm{opt}} = \min\left\{ k \in \{1,\dots,d\}; \mathrm{tr}(S_{Z^\star}) - \sum_{i=1}^{k} \lambda_{Z^\star,i} \leq (d-k) \right\}.$$

Therefore, we can obtain $\widetilde{Z}^\star$ and $\ell^{\mathrm{opt}}$ by solving the eigenvalue equations $S_{Z^\star} u_{Z^\star,i} = \lambda_{Z^\star,i} u_{Z^\star,i}$ for $i = 1,\dots,\ell^{\mathrm{opt}} + 1$.

In practical setting, we set an upper bound parameter $\widetilde{\ell}^{\mathrm{ub}} < \min\{n, d-1\}$ and define $\widetilde{\ell}^{\mathrm{opt}}$ as

$$\widetilde{\ell}^{\mathrm{opt}} = \min\left\{ k \in \{1,\dots,\widetilde{\ell}^{\mathrm{ub}}\}; \mathrm{tr}(S_{Z^\star}) - \sum_{i=1}^{k} \lambda_{Z^\star,i} \leq (d-k) \right\}.$$

Then, we obtain

$$\widetilde{\ell}^{\mathrm{opt}} = \begin{cases} \ell^{\mathrm{opt}}, & \widetilde{\ell}^{\mathrm{ub}} \geq \ell^{\mathrm{opt}}, \\ \widetilde{\ell}^{\mathrm{ub}}, & \widetilde{\ell}^{\mathrm{ub}} < \ell^{\mathrm{opt}}. \end{cases}$$

Therefore, by setting $\widetilde{\ell}^{\mathrm{ub}}$ such that $\widetilde{\ell}^{\mathrm{ub}} \geq \ell^{\mathrm{opt}}$, we do not need to compute the $(\widetilde{\ell}^{\mathrm{ub}} + 1)$th and later eigenvalues. That is, it is sufficient to solve the eigenvalue equation (3) in the procedure II for $i = 1,\dots,\widetilde{\ell}^{\mathrm{ub}}$. As $\ell^{\mathrm{opt}}$ is unknown, we should set a sufficiently large value $\widetilde{\ell}^{\mathrm{ub}}$. Empirically, it may be sufficient to set $\widetilde{\ell}^{\mathrm{ub}} = 1{,}000$ in single-cell sequencing data analysis.

We compare the runtimes and memory usage of RECODE for the number of cells ($n = 1{,}000, 5{,}000, 50{,}000, 100{,}000$) with those of scREOCDE and other imputation methods (Fig ST19). We use the same scRNA-seq data as in the review paper of imputation methods [9], which are Jurkat cell lines [6] ($n = 1{,}000, 5{,}000$) and the bone marrow cells from sample MantonBM6 [14] ($n = 50{,}000, 100{,}000$) created by 10x Genomics. The upper bound parameter $\widetilde{\ell}^{\mathrm{ub}}$ in the fast RECODE algorithm is set to 1,000. The fast algorithm of RECODE (`RECODE_fast`) is faster than the regular algorithm in $n > 1{,}000$ because the fast algorithm works properly when the number of cells is larger than $\widetilde{\ell}^{\mathrm{ub}}$. Compared with that of other imputation methods, the scalability of the fast algorithm of RECODE is superior (4.84). Meanwhile, the memory usage is relatively large because RECODE modifies all entries of the target matrix to real numbers.

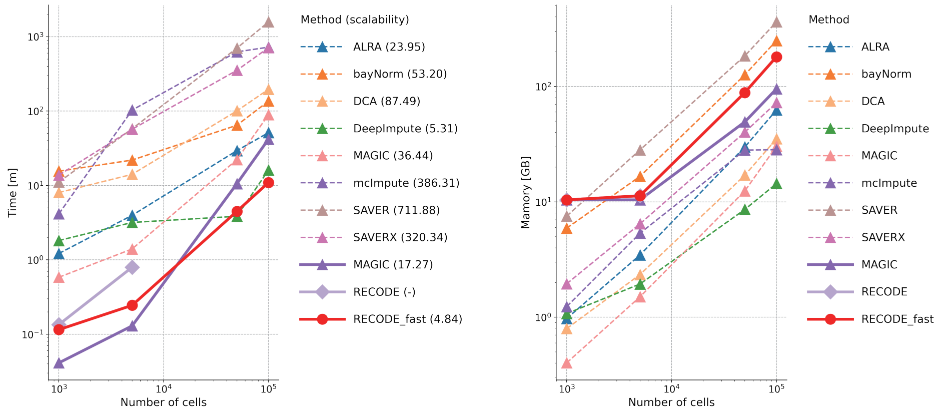


Figure ST19. Comparison of runtime and memory usage of imputation methods and RECODE. The dashed lines are the values of the imputation methods obtained from Hou et al. [9] using the methods computable when $n$ = 100,000. The solid lines are computed using our computing environment (CPU: Intel Xeon W-2223 3.60 GHz). The horizontal axis shows the number of cells ($n = 1{,}000, 5{,}000, 50{,}000, 100{,}000$). The parentheses in the left figure denote the scalability defined by the linear regression with the number of cells. The regular algorithm of RECODE cannot be completed owing to the lack of computational resources when $n = 50{,}000, 100{,}000$.

5.2. **Applicability.** Using biological knowledge, the applicability of RECODE can be examined in advance. From Eq. (20), the empirical NVSN variances $s^2_{Z^\star,i}$ ($i = 1,\ldots,d$), which is given by

$$s^2_{Z^\star,i} = \frac{1}{n-1}\sum_{j=1}^{n}(z^\star_{ij} - \mu_{Z^\star,i})^2$$

where $\mu_{Z^\star,i}$ is the mean of gene $i$ of the normalized data matrix $Z^\star$ and is expected to be greater than 1. Generally, most expressed genes are not directly related to cell identification. In other words, many expressed genes are categorized as non-significant genes in the scRNA-seq data. Therefore, most of them should be distributed around 1.

From this consideration, we classify the scRNA-seq data as follows:

Class A (strongly applicable): Satisfy the following conditions:

(A1) The percentage such that $0 < s^2_{Z^\star,i} < 0.9$ is lower than 1%;

(A2) the peak of the density of $\log(s^2_{Z^\star,i})$ is in the interval $[-0.1, 0.1]$.

Class B (weakly applicable): Satisfy (A1) but do not satisfy (A2).

Class C (inapplicable): Do not satisfy (A1) and (A2).

(A1) indicates that most of the estimated noise variances are not greater than the observed variances. This implies that the observed data contain the estimated noise. (A2) represents the majority of variances $s^2_{Z^\star,i}$ distributed around 1. It corresponds to the aforementioned biological knowledge. Satisfying both (A1) and (A2) indicates good estimations of the noise variances (Class A: strongly applicable). By satisfying (A1) but not (A2), the scRNA-seq data might contain noise other than the estimated noise. In such a case, RECODE may not remove the effects of the additional noises (Class B: weakly applicable). Without satisfying both (A1) and (A2), the modeling of noise may be inappropriate (Class C: inapplicable). The scRNA-seq data used in Section 3.4 are categorized as Class A. Moreover, in the 10X Chromium datasets [6], all scRNA-seq data examined are categorized as Class A (Fig ST20). The applicability to other library creation machines, such as Drop-seq or Smart-seq, is investigated in the main manuscript (see Fig 4 in the main manuscript).

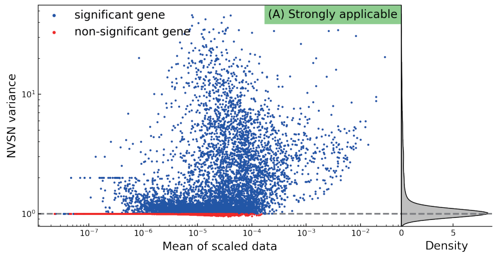


Figure ST20. Applicability of RECODE to scRNA-seq data (3k PBMCs from a Healthy Donor, used in Section 4.3). The left box shows the scatter plots of the mean of the scaled data and the variance of the normalized data. The right side is the density of variance of normalized data.

5.3. **Extension to single-cell epigenome data.** We introduce an extension of RECODE to scATAC-seq data (single-cell assay for transposase-accessible chromatin using sequencing data), which are one of the single-cell epigenome data, to evaluate genome-wide chromatin accessibility. The features of scATAC-seq data denote accessible DNA regions. The number of features of scATAC-seq data is generally larger than that of scRNA-seq data. Therefore, the effect of the COD may be significant in scATAC-seq data analysis.

To apply RECODE to scATAC-seq data, we need to address the error of double counts. As scATAC-seq sequences the subregions of the DNA double helix, it normally counts two times per region. However, it sometimes loses one side of the double helix. Therefore, the scATAC-seq data mainly consist of even counts; however, they sometimes contain odd counts (Fig ST21a left). To modify such errors, we propose the odd–even stabilization:

$$c_{ij}^{\mathsf{stab}} := \lceil c_{ij}/2 \rceil, \quad i = 1, \ldots, d, \ j = 1, \ldots, n.$$

Here, $c_{ij}$ represents the scATAC-seq data with the $i$th peak (feature/subregion) and $j$th sample, and $\lceil\cdot\rceil$ is the ceiling function. After the odd–even stabilization, the frequency of the values in scATAC-seq data smoothly decreases (Fig ST21a right), and the applicability of RECODE becomes strongly applicable (Fig ST21b). Therefore, when applying RECODE to scATAC-seq data, odd–even stabilization is used as the first preprocessing.

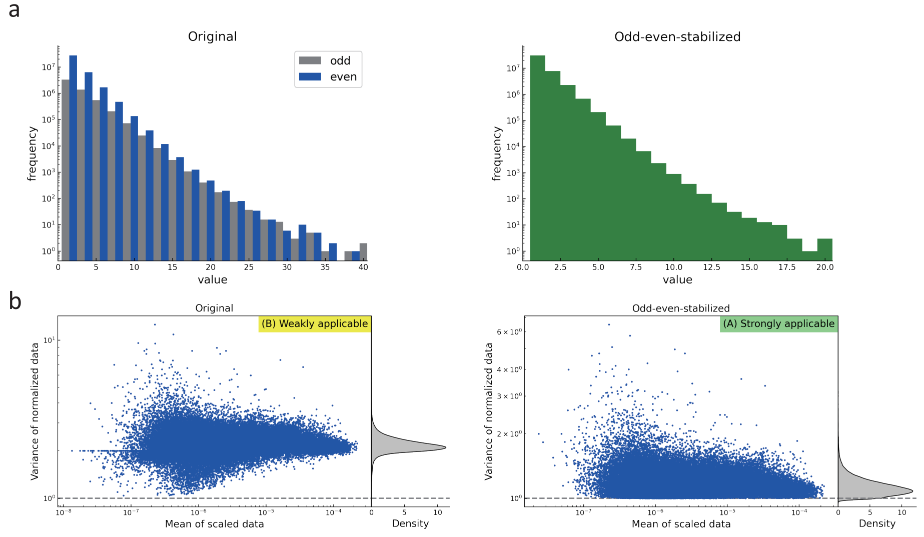


Figure ST21. Odd–even stabilization for scATAC-seq data. **a**, Histogram of count values in original and odd–even stabilized scATAC-seq data. **b**, Applicability RECODE to original and odd–even stabilized scATAC-seq data.

We show the formulation of RECODE for scATAC-seq data as follows.

**RECODE for scATAC-seq data**

For the scATAC-seq data matrix $C = (c_{ij}) \in \mathbb{R}^{d\times n}$ that does not contain features with all zero ($c_{i1} + \cdots + c_{in} \neq 0$ for all $i$), RECODE defines the modified data matrix

$$\widetilde{C} := \widehat{F}^{-1}(U_{\widehat{F}(C)}\widetilde{\Lambda}^{1/2}_{\widehat{F}(C),\ell^{\mathrm{opt}}}\Lambda^{-1/2}_{\widehat{F}(C)}U^{\mathrm{T}}_{\widehat{F}(C)}[\widehat{F}(C) - \overline{\widehat{F}(C)}] + \overline{\widehat{F}(C)}).$$

Here,

$$\widehat{F} : \mathbb{R}^{d\times n} \to \mathbb{R}^{d\times n} \text{ such that } [\widehat{F}(C)]_{ij} := f_{\mathrm{E}(c^{\mathtt{stab}}_{ij}/(t^{\mathtt{stab}}_j)^2)}(c^{\mathtt{stab}}_{ij}/t^{\mathtt{stab}}_j),$$

$$t^{\mathtt{stab}}_j = \sum_{i=1}^{d} c^{\mathtt{stab}}_{ij},$$

$$\ell^{\mathrm{opt}} := \min\left\{k \in \{1,\dots,d\};\ \sum_{i=k+1}^{d} \lambda_{\widehat{F}(C),i} \leq (d-k)\right\}.$$

5.4. **Discussion of extension to future data.** As RECODE does not require assumptions of true data, it has the potential to be applied to other data containing random sampling noises, such as the scATAC-seq data, as mentioned in the previous section. Moreover, the application of RECODE is not restricted to single-cell data. For example, we confirm the strong applicability of RECODE for spatial gene expression data, which are non-single-cell sequencing data (one sample contains few single cells) created by Visium of 10X Genomics, Inc. [6] (Fig ST22). Therefore, in addition to scRNA-seq data, we can expect the application of RECODE to existing or future random sampling data. Furthermore, for non-random sampling data, by analyzing noise variances and applying noise variance-stabilizing normalization in the same manner, RECODE may resolve the COD of such data.

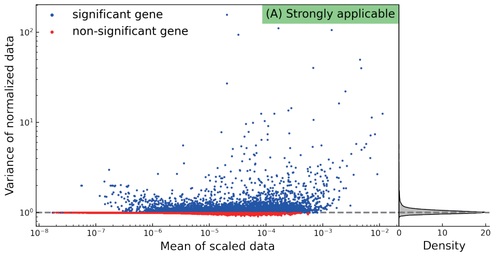


Figure ST22. Applicability of RECODE for spatial gene expression data (non-single-cell data) of normal human prostate (FFPE) generated by Visium of 10X Genomics, Inc. [6].

#### Appendix A. Mathematical notation

In this section, we introduce the mathematical notation used in this paper.

*Bachmann–Landau notation.* For real-valued functions $f(x)$ and $g(x)$, the notation $f(x) = O(g(x))$ as $x \to \infty$ is defined to indicate

$$\limsup_{n\to\infty} \left| \frac{f(x)}{g(x)} \right| < \infty.$$

*Stochastic boundedness.* For a sequence $\{X_d\}$ of random variables and a sequence $\{a_d\}$ of positive numbers, $X_d/a_d$ is said to be *stochastically bounded* as $d \to \infty$, denoted by $X_d = O_{\mathtt{p}}(a_d)$, if the following is satisfied:

$$\forall \epsilon > 0,\ \exists C > 0 \text{ s.t. } \limsup_{d\to\infty} \Pr(|X_d| > Ca_d) < \epsilon.$$

*Convergence in probability.* For a sequence of random variables $\{X_d\}$ and sequences $\{a_d\}$ and $\{b_d\}$ of numbers and positive numbers, respectively, $(X_d - a_d)/b_d$ is said to *converge to zero in probability as* $d \to \infty$, denoted by $X_d = a_d + o_{\mathtt{p}}(b_d)$, if the following is satisfied:

$$\forall \epsilon > 0, \lim_{d\to\infty} \Pr(|X_d - a_d| \geq \epsilon b_d) = 0.$$

When $X_d = Y_d/Z_d$ and $a_d = 1, b_d = 1$ for all $d$, we write

$$Y_d \xrightarrow{\mathtt{p}} Z_d$$

to denote $X_d = 1 + o_{\mathtt{p}}(1)$.

#### Appendix B. Proofs

*Proof of Proposition 1.1.* Based on the assumption regarding the noise terms $e_j$, the difference $e_j - e_{j'}$ for $(j \neq j')$ has a mean of 0 and covariance matrix $2\sigma^2 I$. Therefore, we have

$$\begin{aligned}
\mathrm{E}(\|e_j - e_{j'}\|^2) &= \mathrm{E}\left[\sum_{i=1}^{d} (e_{ij} - e_{ij'})^2\right] \\
&= \sum_{i=1}^{d} \mathrm{E}\left[(e_{ij} - e_{ij'})^2\right] \\
&= \sum_{i=1}^{d} [\mathrm{Var}\,(e_{ij} - e_{ij'}) + \mathrm{E}\,(e_{ij} - e_{ij'})^2] \\
&= 2d\sigma^2.
\end{aligned}$$

Then, we calculate

$$\begin{aligned}
\mathrm{E}(\|x_j - x_{j'}\|^2) &= \mathrm{E}(\|(x_j^{\mathtt{true}} + e_j) - (x_{j'}^{\mathtt{true}} + e_{j'})\|^2) \\
&= \mathrm{E}(\|x_j^{\mathtt{true}} - x_{j'}^{\mathtt{true}}\|^2) + \mathrm{E}(\|e_j - e_{j'}\|^2) \\
&\quad + \mathrm{E}[(x_j^{\mathtt{true}} - x_{j'}^{\mathtt{true}})^{\mathrm{T}}(e_j - e_{j'})] + \mathrm{E}[(e_j - e_{j'})^{\mathrm{T}}(x_j^{\mathtt{true}} - x_{j'}^{\mathtt{true}})] \\
&= \mathrm{E}(\|x_j^{\mathtt{true}} - x_{j'}^{\mathtt{true}}\|^2) + 2\sigma^2 d.
\end{aligned}$$

The last equality follows from the assumption that $x_j^{\mathtt{true}}$, $x_{j'}^{\mathtt{true}}$, $e_j$, and $e_{j'}$ are independent of each other. □

*Proof of Proposition 1.3.* Fix $j, j'$ with $j \neq j'$. By Chebyshev's inequality and condition (2), for any $\epsilon > 0$, we have

$$\Pr(|\|x_j - \mu\|^2 - \mathrm{tr}(S)| \geq \epsilon\, \mathrm{tr}(S)) \leq \frac{\mathrm{Var}(\|x_j - \mu\|^2)}{\epsilon^2 \mathrm{tr}(S)^2} \to 0 \qquad \text{as } d \to \infty,$$

since $\mathrm{E}(\|x_j - \mu\|^2) = \mathrm{tr}(S)$. It follows that

$$\|x_j - \mu\|^2 = \mathrm{tr}(S) + o_{\mathrm{p}}(\mathrm{tr}(S)).$$

Moreover, by Chebyshev's inequality and condition (1), for any $\epsilon > 0$, we have

$$\Pr(|(x_j - \mu)^{\mathrm{T}}(x_{j'} - \mu)| \geq \epsilon\, \mathrm{tr}(S) \leq \frac{\mathrm{tr}(S^2)}{\epsilon^2 \mathrm{tr}(S)^2} \to 0 \qquad \text{as } d \to \infty$$

because $\mathrm{E}((x_j - \mu)^{\mathrm{T}}(x_{j'} - \mu)) = 0$ and $\mathrm{Var}((x_j - \mu)^{\mathrm{T}}(x_{j'} - \mu)) = \mathrm{tr}(S^2)$ under the assumption that $x_j$ and $x_{j'}$ are independent. It follows that

$$(x_j - \mu)^{\mathrm{T}}(x_{j'} - \mu) = o_{\mathrm{p}}(\mathrm{tr}(S)).$$

Note that for any $j \neq j'$,

$$\|x_j - x_{j'}\|^2 = \|x_j - \mu\|^2 + \|x_{j'} - \mu\|^2 - 2(x_j - \mu)^{\mathrm{T}}(x_{j'} - \mu).$$

Thus, we have

$$\|x_j - x_{j'}\|^2 = 2\mathrm{tr}(S) + o_{\mathrm{p}}(\mathrm{tr}(S)).$$

□

*Sketch of Proof of Proposition 1.4.* Under condition (c0), for $i$ such that $\alpha_i \in (0, 1]$, it holds as $d \to \infty$ and $n \to \infty$ that

$$\frac{\lambda_{X,i}}{\lambda_i} = 1 + \frac{O(d)}{n\lambda_i} + o_{\mathrm{p}}(1); \tag{28}$$

see [22, Theorem 3.1 and Theorem 3.3]. If $d^{1-\alpha_i}/n \to 0$, then $O(d)/(n\lambda_i) \to 0$, and the convergence $\lambda_{X,i} \xrightarrow{\mathrm{P}} \lambda_i$ (consistency of eigenvalues) holds. Otherwise, $O(d)/(n\lambda_i)$ remains in the equation (28). This implies the inconsistency of eigenvalues; $\lambda_{X,i} \overset{\mathrm{P}}{\not\to} \lambda_i$. □

*Proof of Lemma 3.1.* Recall that $P$ is an $n \times n$ matrix with all entries $1/n$. As $P^2 = P$, we have $\overline{X}P = \overline{X}$ and $(X - \overline{X})P = O$. Hence,

$$\begin{aligned}\overline{Y} &= YP \\ &= U_X L U_X^{\mathrm{T}}(X - \overline{X})P + \overline{X}P. \\ &= \overline{X}.\end{aligned}$$

Then, $\mu_Y = \mu_X$. Using $\overline{Y} = \overline{X}$ and the eigenvalue decomposition $S_X = U_X \Lambda_X U_X^{\mathrm{T}}$, we have

$$\begin{aligned}S_Y &= \frac{1}{n-1}(Y - \overline{Y})(Y - \overline{Y})^{\mathrm{T}} \\ &= \frac{1}{n-1} U_X L U_X^{\mathrm{T}}(X - \overline{X})(X - \overline{X})^{\mathrm{T}} U_X L U_X^{\mathrm{T}} \\ &= U_X L U_X^{\mathrm{T}} S_X U_X L U_X^{\mathrm{T}} \\ &= U_X L \Lambda_X L U_X^{\mathrm{T}}.\end{aligned}$$

Here, $\Lambda_X \coloneqq \mathrm{diag}(\lambda_{X,1}, \ldots, \lambda_{X,d}) \in \mathbb{R}^{d\times d}$. Therefore, defining $D, U_Y \in \mathbb{R}^{d\times d}$ as

$$D \coloneqq \mathrm{diag}(L_{11}^2 \lambda_{X,1}, \ldots, L_{dd}^2 \lambda_{X,d}),$$
$$U_Y \coloneqq (u_{X,1}, \ldots, u_{X,d}),$$

we obtain the eigenvalue decomposition $S_Y = U_Y D U_Y^{\mathrm{T}}$. □

*Proof of Theorem 3.5.* From Eq. (14), we have

$$\widetilde{x}_j - \widetilde{x}_{j'} = U_X \widetilde{\Lambda}_{X,m}^{1/2} \Lambda_X^{-1/2} U_X^{\mathrm{T}} (x_j - x_{j'}).$$

Then, we have

$$\begin{aligned}
\|\widetilde{x}_j - \widetilde{x}_{j'}\|^2 &= (\widetilde{x}_j - \widetilde{x}_{j'})^{\mathrm{T}} (\widetilde{x}_j - \widetilde{x}_{j'}) \\
&= [U_X \widetilde{\Lambda}_{X,m}^{1/2} \Lambda_X^{-1/2} U_X^{\mathrm{T}} (x_j - x_{j'})]^{\mathrm{T}} [U_X \widetilde{\Lambda}_{X,m}^{1/2} \Lambda_X^{-1/2} U_X^{\mathrm{T}} (x_j - x_{j'})]. \\
&= [(x_j - x_{j'})^{\mathrm{T}} U_X \Lambda_X^{-1/2} \widetilde{\Lambda}_{X,m}^{1/2}][\widetilde{\Lambda}_{X,m}^{1/2} \Lambda_X^{-1/2} U_X^{\mathrm{T}} (x_j - x_{j'})]. \\
&= \|\widetilde{\Lambda}_{X,m}^{1/2} \Lambda_X^{-1/2} U_X^{\mathrm{T}} (x_j - x_{j'})\|^2.
\end{aligned}$$

Since $\widetilde{\Lambda}_{X,m}^{1/2} \Lambda_X^{-1/2}$ is a diagonal matrix and $[\widetilde{\Lambda}_{X,m}^{1/2} \Lambda_X^{-1/2}]_{ii} = 0$ for $i > m$, we have

$$\|\widetilde{x}_j - \widetilde{x}_{j'}\|^2 = \|\widetilde{\Lambda}_{X,m}^{1/2} \Lambda_X^{-1/2} U_X^{\mathrm{T}} (x_j - x_{j'})\|^2 = \|\widehat{\Lambda}_{X,m}^{1/2} U_{X,m}^{\mathrm{T}} (x_j - x_{j'})\|^2,$$

where $\widehat{\Lambda}_{X,m}^{1/2} = \mathrm{diag}(\widetilde{\lambda}_{X,1}^{1/2} \lambda_{X,1}^{-1/2}, \ldots, \widetilde{\lambda}_{X,m}^{1/2} \lambda_{X,m}^{-1/2}) \in \mathbb{R}^{m\times m}$ and $U_{X,m} = (u_{X,1} \ldots u_{X,m}) \in \mathbb{R}^{d\times m}$. As $\|Lb\|^2 \le \max_i \{L_{ii}^2\} \|b\|^2$ holds for a diagonal matrix $L$ and vector $b$ and because $\widetilde{\lambda}_{X,i} \le \lambda_{X,i}$ for $i = 1, \ldots, d$, we have

$$\begin{aligned}
\|\widetilde{x}_j - \widetilde{x}_{j'}\|^2 &\le \max_{i=1,\ldots,m} \{\widetilde{\lambda}_{X,i} / \lambda_{X,i}\} \|U_{X,m}^{\mathrm{T}} (x_j - x_{j'})\|^2 \\
&\le \|U_{X,m}^{\mathrm{T}} (x_j - x_{j'})\|^2.
\end{aligned}$$

Moreover, we have

$$\begin{aligned}
\|U_{X,m}^{\mathrm{T}} (x_j - x_{j'})\|^2 &= \|U_{X,m}^{\mathrm{T}} (x_j^{\mathtt{true}} - x_{j'}^{\mathtt{true}}) + U_{X,m}^{\mathrm{T}} (e_j - e_{j'})\|^2 \\
&= \|U_{X,m}^{\mathrm{T}} (x_j^{\mathtt{true}} - x_{j'}^{\mathtt{true}})\|^2 + \|U_{X,m}^{\mathrm{T}} (e_j - e_{j'})\|^2 \\
&\quad + (e_j - e_{j'})^{\mathrm{T}} U_{X,m} U_{X,m}^{\mathrm{T}} (x_j^{\mathtt{true}} - x_{j'}^{\mathtt{true}}) \\
&\quad + (x_j^{\mathtt{true}} - x_{j'}^{\mathtt{true}})^{\mathrm{T}} U_{X,m} U_{X,m}^{\mathrm{T}} (e_j - e_{j'}).
\end{aligned}$$

From condition (12), for $k \in \{j, j'\}$, we have

$$\begin{aligned}
x_k^{\mathtt{true}} - \mu_{X^{\mathtt{true}}} &= U_X U_X^{\mathrm{T}} (x_k^{\mathtt{true}} - \mu_{X^{\mathtt{true}}}) \\
&= U_X [(u_{X,1} \ldots u_{X,m}\ 0_d \ldots 0_d) + (0_d \ldots 0_d\ u_{X,m+1} \ldots u_{X,d})]^{\mathrm{T}} (x_k^{\mathtt{true}} - \mu_{X^{\mathtt{true}}}) \\
&= U_X (u_{X,1} \ldots u_{X,m}\ 0_d \ldots 0_d)^{\mathrm{T}} (x_k^{\mathtt{true}} - \mu_{X^{\mathtt{true}}}) \\
&= (u_{X,1} \ldots u_{X,m}\ 0_d \ldots 0_d)(u_{X,1} \ldots u_{X,m}\ 0_d \ldots 0_d)^{\mathrm{T}} (x_k^{\mathtt{true}} - \mu_{X^{\mathtt{true}}}) \\
&= U_{X,m} U_{X,m}^{\mathrm{T}} (x_k^{\mathtt{true}} - \mu_{X^{\mathtt{true}}}),
\end{aligned}$$

where $0_d$ is the $d$-dimensional zero vector. Then, we have

$$\begin{aligned}
\|U_{X,m}^{\mathrm{T}} (x_j^{\mathtt{true}} - x_{j'}^{\mathtt{true}})\|^2 &= (x_j^{\mathtt{true}} - x_{j'}^{\mathtt{true}})^{\mathrm{T}} U_{X,m} U_{X,m}^{\mathrm{T}} (x_j^{\mathtt{true}} - x_{j'}^{\mathtt{true}}) \\
&= (x_j^{\mathtt{true}} - x_{j'}^{\mathtt{true}})^{\mathrm{T}} U_{X,m} U_{X,m}^{\mathrm{T}} [(x_j^{\mathtt{true}} - \mu_{X^{\mathtt{true}}}) - (x_{j'}^{\mathtt{true}} - \mu_{X^{\mathtt{true}}})] \\
&= (x_j^{\mathtt{true}} - x_{j'}^{\mathtt{true}})^{\mathrm{T}} [(x_j^{\mathtt{true}} - \mu_{X^{\mathtt{true}}}) - (x_{j'}^{\mathtt{true}} - \mu_{X^{\mathtt{true}}})] \\
&= \|x_j^{\mathtt{true}} - x_{j'}^{\mathtt{true}}\|^2.
\end{aligned}$$

Moreover, from the independence of the noise and true data, we have

$$\begin{aligned}\mathrm{E}(\|\widetilde{x}_j - \widetilde{x}_{j'}\|^2) &\leq \mathrm{E}(\|x_j^{\mathtt{true}} - x_{j'}^{\mathtt{true}}\|^2) + \mathrm{E}(\|U_{X,m}^{\mathrm{T}}(e_j - e_{j'})\|^2)\\ &\quad + \mathrm{E}[(e_j - e_{j'})^{\mathrm{T}} U_{X,m} U_{X,m}^{\mathrm{T}}(x_j^{\mathtt{true}} - x_{j'}^{\mathtt{true}})]\\ &\quad + \mathrm{E}[(x_j^{\mathtt{true}} - x_{j'}^{\mathtt{true}})^{\mathrm{T}} U_{X,m} U_{X,m}^{\mathrm{T}}(e_j - e_{j'})]\\ &= \mathrm{E}(\|x_j^{\mathtt{true}} - x_{j'}^{\mathtt{true}}\|^2) + \mathrm{E}(\|U_{X,m}^{\mathrm{T}}(e_j - e_{j'})\|^2).\end{aligned}$$

From the independence of noise and $e_{kj} - e_{kj'}$ following a distribution with mean 0 and variance $2\sigma^2$, we have

$$\begin{aligned}\mathrm{E}(\|U_{X,m}^{\mathrm{T}}(e_j - e_{j'})\|^2) &= \sum_{k=1}^{d}\sum_{s=1}^{m} u_{X,ks}^2 \mathrm{E}((e_{kj} - e_{kj'})^2)\\ &= \sum_{k=1}^{d}\sum_{s=1}^{m} u_{X,ks}^2 \mathrm{Var}(e_{kj} - e_{kj'})\\ &= 2\sigma^2 \sum_{k=1}^{d}\sum_{s=1}^{m} u_{X,ks}^2\\ &= 2\sigma^2 m.\end{aligned}$$

Here, $u_{X,i} = (u_{X,1i}, \ldots, u_{X,di})^{\mathrm{T}}$. Therefore, we finally obtain

$$\mathrm{E}(\|\widetilde{x}_j - \widetilde{x}_{j'}\|^2) \leq \mathrm{E}(\|x_j^{\mathtt{true}} - x_{j'}^{\mathtt{true}}\|^2) + 2\sigma^2 m.$$

□

*Proof of Theorem 3.6.* From the eigenvalue equation (3) and condition (C1), for $i = m+1, \ldots, d$, we have

$$\begin{aligned}\lambda_{X,i} &= u_{X,i}^{\mathrm{T}} S_X u_{X,i}\\ &= \frac{1}{n-1} u_{X,i}^{\mathrm{T}} (X - \overline{X})(X - \overline{X})^{\mathrm{T}} u_{X,i}\\ &= \frac{1}{n-1} u_{X,i}^{\mathrm{T}} \left[(X^{\mathtt{true}} - \overline{X^{\mathtt{true}}}) + (E - \overline{E})\right]\left[(X^{\mathtt{true}} - \overline{X^{\mathtt{true}}}) + (E - \overline{E})\right]^{\mathrm{T}} u_{X,i}\\ &= \frac{1}{n-1} u_{X,i}^{\mathrm{T}} (E - \overline{E})(E - \overline{E})^{\mathrm{T}} u_{X,i}\\ &= u_{X,i}^{\mathrm{T}} S_E u_{X,i}\,.\end{aligned}$$

□

*Proof of Theorem 4.1.* We fix a gene $i$ and consider the distributions of $x_{ij} = c_{ij}/t_j$ for $j = 1, \ldots, n$. It follows from the conditional probability distribution (18) that

$$\begin{aligned}&\Pr(c_{ij} = k, t_j = t, x_{ij}^{\mathtt{true}} = y)\\ &\quad = \Pr(c_{ij} = k | t_j = t, x_{ij}^{\mathtt{true}} = y)\Pr(t_j = t, x_{ij}^{\mathtt{true}} = y)\\ &\quad = \frac{(ty)^k e^{-ty}}{k!}\Pr(t_j = t, x_{ij}^{\mathtt{true}} = y).\end{aligned}$$

As $\mathrm{E}(Y) = \mathrm{Var}(Y) = \lambda$ for $Y$ following a Poisson distribution with parameter $\lambda$, the mean and variance of $x_{ij} = c_{ij}/t_j$ can be calculated as follows:

$$\begin{aligned}
\mathrm{E}(x_{ij}) &= \sum_y \sum_t \sum_k \frac{k}{t} \Pr(c_{ij} = k, t_j = t, x_{ij}^{\texttt{true}} = y) \\
&= \sum_y \sum_t \sum_k \frac{k}{t} \frac{(ty)^k e^{-ty}}{k!} \Pr(t_j = t, x_{ij}^{\texttt{true}} = y) \\
&= \sum_y \sum_t y \Pr(t_j = t, x_{ij}^{\texttt{true}} = y) \\
&= \mathrm{E}(x_{ij}^{\texttt{true}}),
\end{aligned}$$

$$\begin{aligned}
\mathrm{E}(x_{ij}^2) &= \sum_y \sum_t \sum_k \left(\frac{k}{t}\right)^2 \frac{(ty)^k e^{-ty}}{k!} \Pr(t_j = t, x_{ij}^{\texttt{true}} = y) \\
&= \sum_y \sum_t \frac{(ty)^2 + ty}{t^2} \Pr(t_j = t, x_{ij}^{\texttt{true}} = y) \\
&= \mathrm{E}[(x_{ij}^{\texttt{true}})^2] + \mathrm{E}(x_{ij}^{\texttt{true}}/t_j),
\end{aligned}$$

$$\begin{aligned}
\mathrm{Var}(x_{ij}) &= \mathrm{E}(x_{ij}^2) - \mathrm{E}(x_{ij})^2 \\
&= \mathrm{E}((x_{ij}^{\texttt{true}})^2) + \mathrm{E}(x_{ij}^{\texttt{true}}/t_j) - \mathrm{E}(x_{ij}^{\texttt{true}})^2 \\
&= \mathrm{Var}(x_{ij}^{\texttt{true}}) + \mathrm{E}(x_{ij}^{\texttt{true}}/t_j).
\end{aligned}$$

Since

$$\begin{aligned}
\mathrm{E}(x_{ij}/t_j) &= \sum_y \sum_t \sum_k \frac{k}{t^2} \frac{(ty)^k e^{-ty}}{k!} \Pr(t_j = t, x_{ij}^{\texttt{true}} = y) \\
&= \sum_y \sum_t \frac{y}{t} \Pr(t_j = t, x_{ij}^{\texttt{true}} = y) \\
&= \mathrm{E}(x_{ij}^{\texttt{true}}/t_j),
\end{aligned}$$

we have

$$\mathrm{Var}(x_{ij}) = \mathrm{Var}(x_{ij}^{\texttt{true}}) + \mathrm{E}(x_{ij}/t_j).$$

□

[4] Makoto Aoshima and Kazuyoshi Yata. 高次元の統計学 (統計学 *One Point*) [*High-Dimensional Statistics* (*Statistics One Point*)] (*in Japanese*). Kyoritsu Shuppan, 2019.

[5] Bo Ding, Lina Zheng, Yun Zhu, Nan Li, Haiyang Jia, Rizi Ai, Andre Wildberg, and Wei Wang. Normalization and noise reduction for single cell RNA-seq experiments. *Bioinformatics*, 31(13):2225–2227, 2015.

[6] 10X Genomics. Single cell gene expression datasets. https://support.10xgenomics.com/single-cell-gene-expression/datasets. (accessed 31 Mar. 2022).

[7] Dominic Grün, Lennart Kester, and Alexander Van Oudenaarden. Validation of noise models for single-cell transcriptomics. *Nature methods*, 11(6):637–640, 2014.
[8] Peter Hall, James Stephen Marron, and Amnon Neeman. Geometric representation of high dimension, low sample size data. *Journal of the Royal Statistical Society: Series B (Statistical Methodology)*, 67(3):427–444, 2005.
[9] Wenpin Hou, Zhicheng Ji, Hongkai Ji, and Stephanie C. Hicks. A systematic evaluation of single-cell rna-sequencing imputation methods. *Genome Biology*, 21(1), 2020.
[10] Saiful Islam, Amit Zeisel, Simon Joost, Gioele La Manno, Pawel Zajac, Maria Kasper, Peter Lönnerberg, and Sten Linnarsson. Quantitative single-cell RNA-seq with unique molecular identifiers. *Nature methods*, 11(2):163, 2014.
[11] Gioele La Manno, Ruslan Soldatov, Amit Zeisel, Emelie Braun, Hannah Hochgerner, Viktor Petukhov, Katja Lidschreiber, Maria E Kastriti, Peter Lönnerberg, Alessandro Furlan, et al. RNA velocity of single cells. *Nature*, 560(7719):494–498, 2018.
[12] Laurens van der Maaten and Geoffrey Hinton. Visualizing data using t-SNE. *Journal of machine learning research*, 9(Nov):2579–2605, 2008.
[13] Leland McInnes, John Healy, and James Melville. Umap: Uniform manifold approximation and projection for dimension reduction. *arXiv preprint arXiv:1802.03426*, 2018.
[14] Aviv Regev. Census of immune cells in human cell atlas data portal. https://data.humancellatlas.org/explore/projects/cc95ff89-2e68-4a08-a234-480eca21ce79. (accessed 31 Mar. 2022).
[15] John G Saw, Mark CK Yang, and Tse Chin Mo. Chebyshev inequality with estimated mean and variance. *The American Statistician*, 38(2):130–132, 1984.
[16] Hideaki Shimazaki and Shigeru Shinomoto. A method for selecting the bin size of a time histogram. *Neural computation*, 19(6):1503–1527, 2007.
[17] Warren S Torgerson. Multidimensional scaling: I. theory and method. *Psychometrika*, 17(4):401–419, 1952.
[18] Florian Wagner, Dalia Barkley, and Itai Yanai. Accurate denoising of single-cell RNA-Seq data using unbiased principal component analysis. 2019.
[19] Kazuyoshi Yata and Makoto Aoshima. PCA consistency for non-Gaussian data in high dimension, low sample size context. *Communications in Statistics—Theory and Methods*, 38(16-17):2634–2652, 2009.
[20] Kazuyoshi Yata and Makoto Aoshima. Effective PCA for high-dimension, low-sample-size data with singular value decomposition of cross data matrix. *Journal of Multivariate Analysis*, 101(9):2060–2077, 2010.
[21] Kazuyoshi Yata and Makoto Aoshima. Effective PCA for high-dimension, low-sample-size data with noise reduction via geometric representations. *Journal of Multivariate Analysis*, 105(1):193–215, 2012.
[22] Kazuyoshi Yata and Makoto Aoshima. PCA consistency for the power spiked model in high-dimensional settings. *Journal of multivariate analysis*, 122:334–354, 2013.

[1]Institute for the Advanced Study of Human Biology, Kyoto University Institute for Advanced Study, Kyoto University, Yoshida-Konoe-cho, Sakyo-ku, Kyoto 606-8501, Japan.

[2]Department of Anatomy and Cell Biology, Graduate School of Medicine, Kyoto University, Yoshida-Konoe-cho, Sakyo-ku, Kyoto 606-8501, Japan.

[3]The Hakubi Center for Advanced Research, Kyoto University, Yoshida-Konoe-cho, Sakyo-ku, Kyoto 606-8501, Japan.

[4]Graduate School of Human Development and Environment, Kobe University, 3-11 Tsurukabuto, Nada-ku, Kobe 657-8501 Japan.

[5]Center for Advanced Intelligence Project, RIKEN. Nihonbashi 1-chome Mitsui Building, 15th floor, 1-4-1 Nihonbashi, Chuo-ku, Tokyo 103-0027, Japan

[6]Center for iPS Cell Research and Application, Kyoto University, 53 Kawahara-cho, Shogoin, Sakyo-ku, Kyoto 606-8507, Japan.

[7]Center for Advanced Study, Kyoto University Institute for Advanced Study, Kyoto University.
